# Cytoplasmic CaMKIIδ-B prevents myocardial recovery in heart failure

**DOI:** 10.1101/2025.05.16.654509

**Authors:** Thomas G. Martin, Dakota R. Hunt, Christopher C. Ebmeier, Abhishek P. Dhand, Christina Alamana, Joseph C. Cleveland, Sharon L. Graw, Sarah Bruner, Michael R. Bristow, Luisa Mestroni, Matthew R.G. Taylor, Jason A. Burdick, Amrut V. Ambardekar, Peter M. Buttrick, Leslie A. Leinwand

## Abstract

Restoration of cardiac function in patients with advanced heart failure is rare, and the molecular processes that regulate recovery are unknown. To identify potential mechanisms, we studied paired myocardial samples before and after left ventricular assist device therapy, where significant cardiac functional recovery occurred in ∼25% of patients. We found that expression of the nuclear B isoform of Ca^2+^/calmodulin-dependent protein kinase IIδ (CaMKIIδ-B) inversely correlated with recovery. Furthermore, increased phosphorylation near the CaMKIIδ-B nuclear localization signal in non-responders prevented its auto-activation dependent nuclear translocation. Expression of a cytoplasm-restricted CaMKIIδ-B in cardiomyocytes dramatically remodeled the phospho-proteome and impaired contractility, while a nuclear-competent version did not. Modulating CaMKIIδ subcellular localization may thus represent a therapeutic strategy for advanced heart failure.

## Introduction

Heart failure is the leading cause of morbidity and mortality in the developed world (*1*); however, most current therapies treat heart failure symptoms and do not target the underlying molecular causes of the disease (*2*–*4*). Thus, while patients with heart failure are living longer than ever before, their quality of life remains poor and the burden of heart failure on the healthcare system is increasing (*1, 5, 6*). The only current ‘cure’ for heart failure is a heart transplant, access to which is limited due to a shortage of healthy donor hearts (*7*). Notably, substantial clinical evidence indicates that a small proportion of patients treated with guideline-directed therapies experience recovery from heart failure, characterized by reduced left ventricular (LV) dilatation and improved LV systolic function (i.e., reverse remodeling) (*8*). This reverse remodeling is associated with dramatic improvements in patient survival and quality of life (*8*–*10*). However, despite considerable interest from both clinical and basic science perspectives in identifying therapeutic targets for myocardial recovery, the molecular mechanisms that regulate this process remain largely unknown.

Left ventricular assist device (LVAD) therapy, which mechanically unloads the LV and takes over cardiac circulatory function (*11*), is the treatment that is most often associated with reverse remodeling and recovery (*8, 12*–*14*). LVAD therapy also necessitates the removal of some LV myocardium during device placement, thus generating a pre-treatment sample for future paired comparisons. A recent investigation into the molecular biology of recovery by bulk RNA-sequencing found few pre-LVAD gene expression differences between patients who went on to recover and those who did not (*13*), indicating a striking transcriptional similarity between response subgroups before treatment. This finding suggested that molecular features of a favorable response might only manifest after treatment (i.e., they are induced by LVAD). In support of this, single-nucleus sequencing of pre- and post-LVAD myocardium from responders and non-responders identified cell type-specific recovery signatures post-LVAD, where decreased expression of pro-inflammatory genes in macrophages and fibroblasts was the most discriminating feature (*14*). However, as is true with inflammation and heart failure (*15*), whether changes in pro-inflammatory gene expression are a cause or consequence of recovery is not clear. Further, this previous study found that, despite improved gross cardiac contractility denoted by increased systolic function, cardiomyocytes did not revert to a ‘healthy’ transcriptional state in recovery (*14*). These findings support the conclusion that gene expression changes alone do not explain the differential recovery responses. Further, they suggest that other molecular factors contribute to improved cardiac functional outcomes in the subset of patients who recover from advanced heart failure.

## Results

### Transcriptome remodeling with mechanical circulatory support

Among 41 available paired pre- and post-LVAD patient samples (**Data S1**), we identified 10 responders and 9 non-responders based on LV structural and functional outcomes (see methods) (**Fig. 1A-B**). All patients had non-ischemic dilated cardiomyopathy and received LVAD as a bridge-to-transplant. Both outcome groups displayed similar age ranges, medication history, and clinical characteristics before starting therapy (**Table S1-S2**). To examine transcriptional differences, we performed bulk RNA-sequencing to assess gene expression differences in heart failure (pre-LVAD) and post-LVAD samples compared to non-failing controls. We found that mechanical unloading partially restored transcriptional features of heart failure to non-failing control levels (**Fig. S1)**, including reversing pro-inflammatory gene expression (**Fig. S1I**). However, few genes were significantly differentially expressed between responders and non-responders pre-LVAD, in agreement with previous work (*13*), or post-LVAD (**Fig. S2**). Although total gene expression levels were not different between groups, we reasoned that a paired analysis of gene expression changes from pre-to post-LVAD might reveal transcriptional signatures of recovery. We therefore analyzed the RNA-seq data for differences in LVAD treatment effects between responders and non-responders. Gene set enrichment analysis revealed that increased RNA splicing factor gene expression was positively associated with recovery (**Fig. 1C**), while genes involved in the immune response were negatively associated (**Fig. 1D**).

**Fig. 1.**
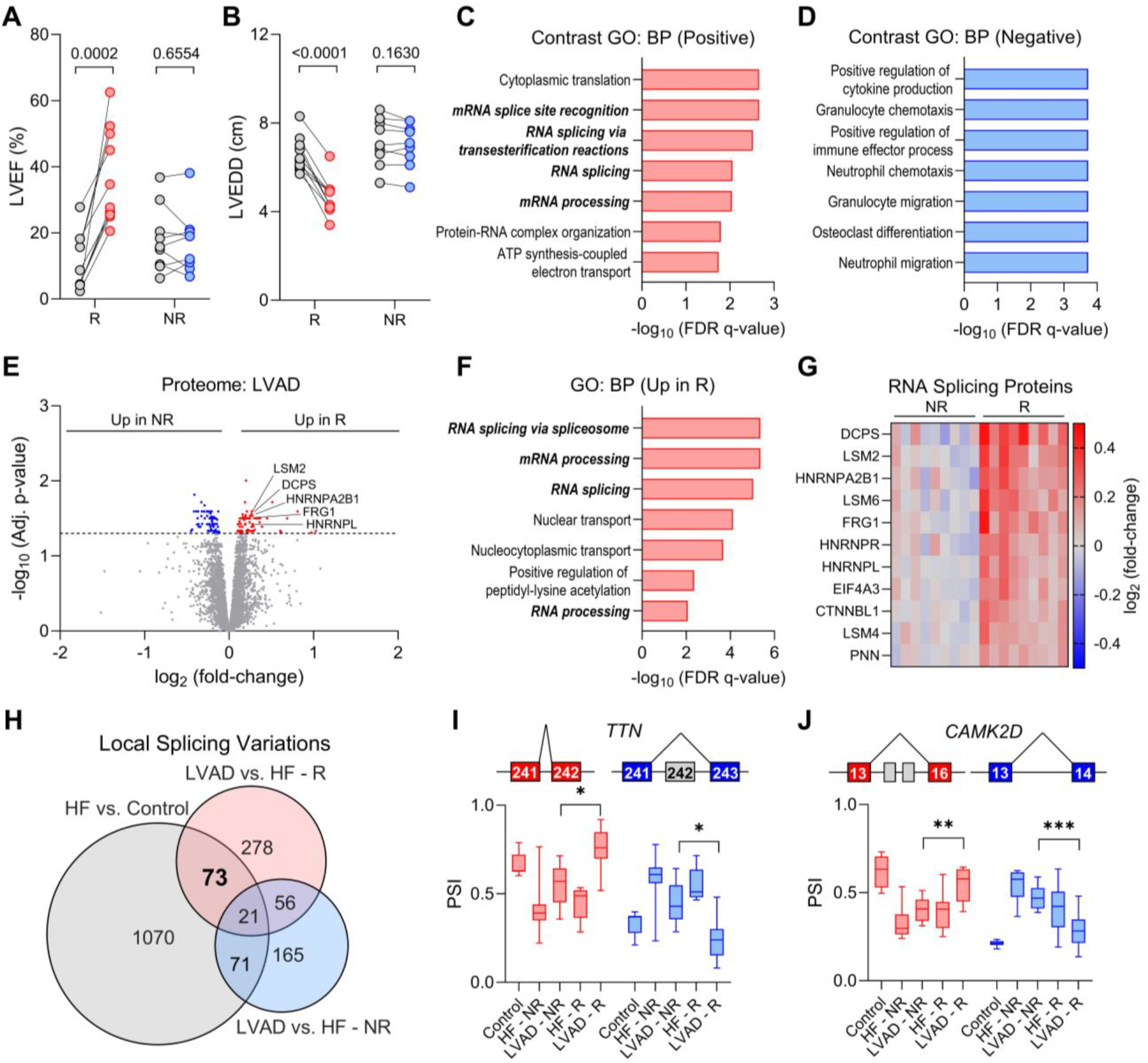
Myocardial recovery in heart failure is associated with differential alternative splicing. **(A-B)** Left ventricular ejection fraction (LVEF) (A) and LV end diastolic dimension (LVEDD) (B) in pre- and post-LVAD in responders (R) and non-responders (NR); n = 10 R, 9 NR; paired two-tailed t-test. **(C-D)** GSEA gene ontology (GO) Biological Process (BP) analysis of bulk RNA-seq data for genes with expression changes pre-to post-LVAD that were positively (C) or negatively (D) associated with a favorable response to LVAD. **(E)** Volcano plot displaying quantitative proteomics results from post-LVAD responder (R) and non-responder (NR) myocardium; n = 9/group. **(F)** Gene ontology (GO) Biological Process pathway enrichment analysis of the significantly upregulated proteins in Responders. **(G)** Heat map displaying the significantly differentially expressed RNA splicing proteins between post-LVAD R and NR. **(H)** Venn diagram displaying the number of local splicing variations identified between HF (n = 19) and control (n = 5) by RNA-seq and the differential alternative splicing responses post-LVAD in R (n = 10) and NR (n = 9) patients. **(I)** *TTN* exon 242 alternative splicing map and quantification between groups. **(J)** *CAMK2D* exon 16 and 14 alternative splicing maps and quantification between groups; PSI = proportion spliced in; n = 5 control, 9 HF-NR and LVAD-NR, 10 HF-R and LVAD-R; *p < 0.05, **p < 0.01, ***p < 0.001 by an independent two-sample Mann-Whitney U-test (Wilcoxon test in MAJIQ).

### Unique RNA alternative splicing changes with heart failure recovery

Over 90% of the ∼20,000 protein-coding genes in the human genome undergo alternative splicing, increasing the complexity of the transcriptome by an order of magnitude (*16, 17*). The generation of multiple transcripts from a single gene increases protein diversity and can thus modulate cellular function. To examine whether the transcriptional differences in splicing factor expression translated to meaningful differences at the protein level, we performed tandem-mass-tag (TMT) quantitative proteomics on post-LVAD responder and non-responder samples. This approach identified 154 differentially expressed proteins between groups, including increased expression of RNA splicing factors in responders (**Fig. 1E-G, Fig. S3-4**). Given this support for the role of alternative splicing in heart failure development and recovery, we next mined the RNA-seq dataset to detect and quantify local splicing variations (LSVs) (*18*). We identified 1,235 LSVs that were significantly impacted in heart failure versus control samples and 411 that changed with mechanical unloading (**Fig. S5-7**). Among the heart failure-associated splicing changes, 34% of altered cassette exon (i.e., exon-skipping) events were predicted to affect protein domains involved in cellular signal transduction (**Fig. S8-9, Table S3**), suggesting that alternative splicing contributes to phenotypic changes in heart failure. To determine if LVAD induced unique alternative splicing changes in heart failure patients who recovered function, we examined LSVs in the responder and non-responder groups compared to heart failure. This analysis revealed that responders and non-responders had distinct alternative splicing responses to LVAD, with 351 and 236 LSVs unique to each group, respectively (**Fig. 1H**).

Among 73 heart failure-associated LSVs that were modified in responders, one notable example was exon 242 inclusion in *TTN* (**Fig. 1I**). *TTN* encodes the giant protein titin, which regulates myofibril passive tension (*19*). Exon 242 encodes an 89 amino acid immunoglobulin (Ig) domain that regulates protein-protein interactions (**Fig. S9**) in the I-band region of titin, which is the primary determinant of titin’s elastic properties (*20*). Thus, alternative splicing in this region is expected to impact sarcomere protein interactions and titin-based myofibril stiffness. Our follow-up analysis of this event using exon-specific qPCR revealed that, while exon 242 inclusion increased across all patients after LVAD compared with heart failure, variability between responders and non-responders contributed to a modest, but non-significant change between these groups (**Fig. S10**).

### Myocardial recovery coincides with decreased CAMK2D exon 14 inclusion

Among the most significant recovery-associated splicing changes identified in the RNA-seq dataset were increased exon 16 and decreased exon 14 inclusion in *CAMK2D* (**Fig. 1J**), which encodes Ca^2+^/calmodulin-dependent protein kinase IIδ (CaMKIIδ). CaMKIIδ is a serine/threonine kinase that regulates contractility and immune response signaling in cardiomyocytes and its chronic hyperactivation is implicated in multiple forms of heart disease (*21*–*23*). Alternative splicing of *CAMK2D* exons 14-16 generates four cardiac isoforms (**Fig. 2A**), which differ in their subcellular localization (*24, 25*). Isoform-specific qPCR analysis corroborated the RNA-seq finding that the δ-B isoform, which includes a nuclear localization signal (NLS) encoded by exon 14, increased in heart failure (**Fig. S11**). Notably, exon 14 inclusion was increased in non-responder patients both pre- and post-LVAD (**Fig. 2B**) and displayed a strong inverse correlation with LV functional and structural outcomes (**Fig. 2C-D**). Targeted RT-PCR and qPCR analyses further validated that there was a shift in the dominant *CAMK2D* isoform from δ-9 (exons 13-16-17) in non-failing controls to δ-B (exons 13-14-17) in heart failure, which was reversed only in patients who experienced functional recovery after LVAD (**Fig. 2E-G**).

**Fig. 2.**
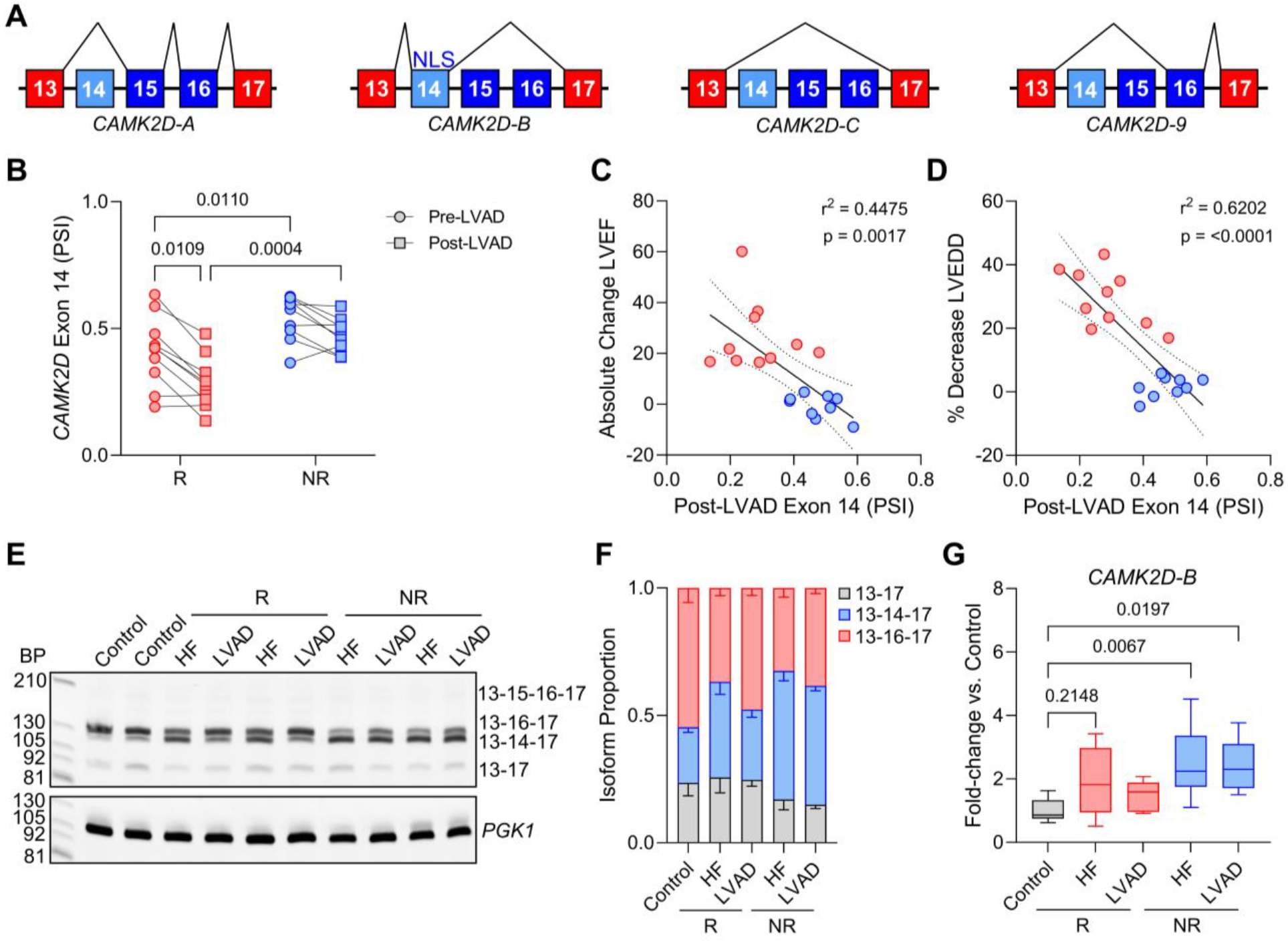
Increased *CAMK2D* exon 14 inclusion in heart failure is reversed in recovery. **(A)** Graphical representation of cardiac *CAMK2D* isoforms and their respective variable region (exons 13-17) alternative splicing; NLS = nuclear localization signal. **(B)** *CAMK2D* exon 14 proportion spliced in (PSI) in pre- and post-LVAD responders and non-responders from the RNA-seq data; two-way ANOVA with Tukey’s post-hoc test. **(C-D)** Linear regression analysis of exon 14 PSI (post-LVAD) vs. absolute change in LVEF (B) and % change in LVEDD (C) on LVAD therapy; confidence interval = 95%. **(E)** Representative RT-PCR acrylamide gel for *CAMK2D* cardiac isoforms and *PGK1* housekeeping gene in non-failing controls, HF, and post-LVAD R and NR; BP = base-pairs. **(F)** Proportion *CAMK2D* isoform expression from the RT-PCR; n = 6 Control, 10 HF and LVAD R, 9 HF and LVAD NR. **(G)** Quantitative RT-PCR analysis of *CAMK2D-B* in Control, HF (pre-LVAD), and post-LVAD normalized to the housekeeping gene *PGK1* and plotted as a fold-change vs. non-failing control expression; n = 10 R, 9 NR; one-way ANOVA with Tukey’s post-hoc test.

### CaMKIIδ NLS phosphorylation status predicts recovery

To identify potential additional molecular factors regulated in recovery, we next performed TMT quantitative phospho-proteomics on the post-LVAD responder and non-responder samples. Remarkably, the most significantly differentially phosphorylated sites identified were three serine residues in CaMKIIδ (S332, S333, and S334), which increased in non-responders (**Fig. 3A-B**). These sites are located immediately downstream of the NLS encoded by exon 14 (**Fig. 3A**) and their phosphorylation was previously implicated in regulating nuclear localization of CaMKIIδ (*26*–*28*). Western blot analyses did not identify differences in CaMKIIδ regulatory domain M281/282 oxidation or T287 autophosphorylation between groups (**Fig. 3C-E**), suggesting similar kinase activation status between responders and non-responders (*29*). However, S332 phosphorylation increased in heart failure and then was fully restored to non-failing control levels in patients who experienced functional recovery (**Fig. 3C, 3F**). In non-responders, pre-LVAD p-S332 levels were ∼2-fold higher than in responders and remained elevated following mechanical unloading (**Fig. 3C, 3F**). Pre- and post-LVAD S332 phosphorylation displayed a significant inverse correlation with functional recovery (**Fig. S12, 3G**), supporting the potential utility of this event as a biomarker to predict which patients are poised for heart failure recovery with LVAD.

**Fig. 3.**
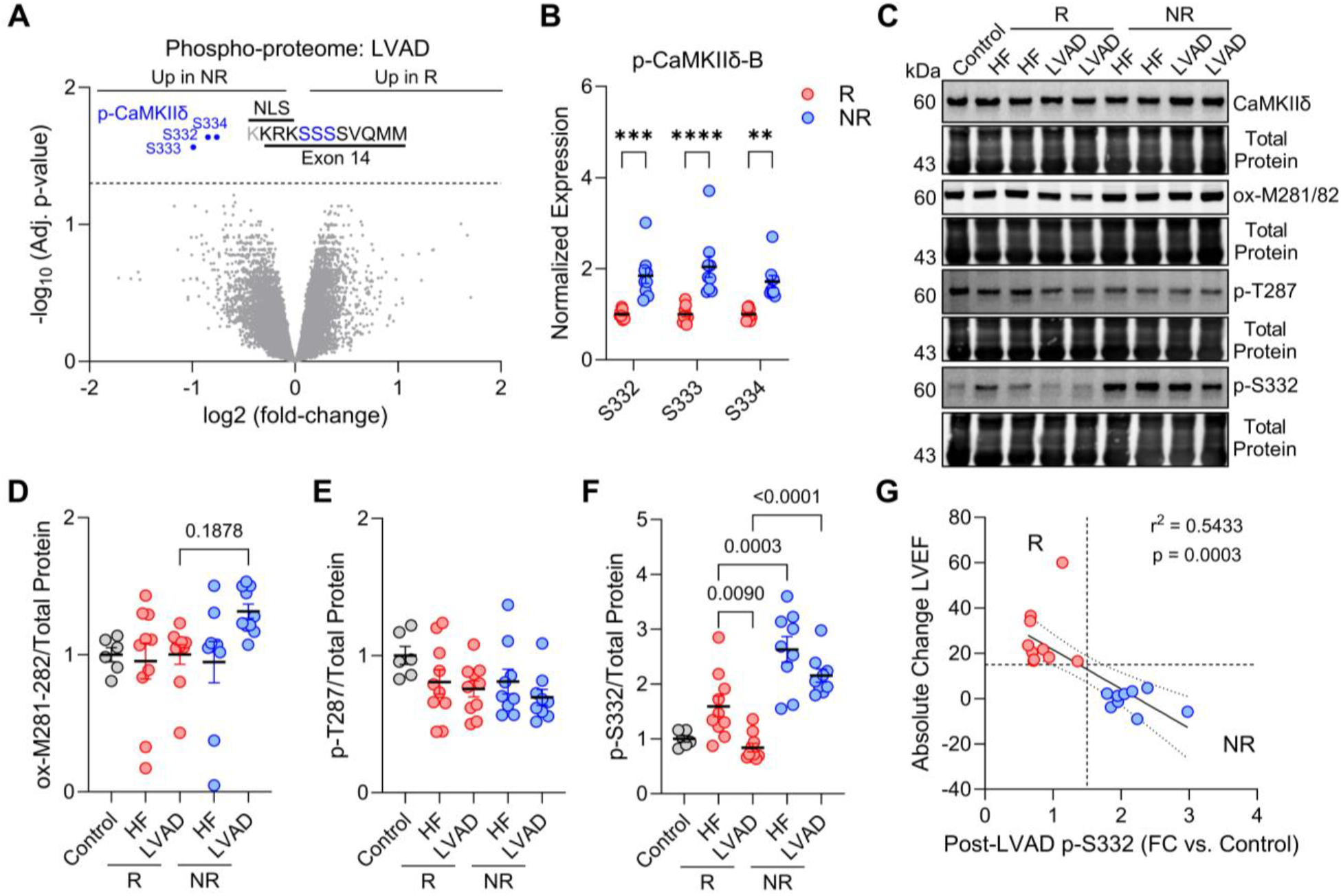
CaMKIIδ-B S332-S334 phosphorylation predicts poor functional recovery. **(A)** Volcano plot displaying phospho-proteomics results from post-LVAD responder (R) and non-responder (NR) myocardium; n = 9/group. **(B)** CaMKIIδ-B S332, S333, and S334 phosphorylation between R and NR post-LVAD; two-tailed t-test; *p < 0.05, **p < 0.01, ****p < 0.0001. **(C)** Representative western blots for total CaMKIIδ, ox-M281/282, p-T287, and p-S332 in non-failing control, pre-LVAD (HF) R and NR, and post-LVAD R and NR. **(D-F)** Ox-M281/282 CaMKIIδ (D), p-T287 CaMKIIδ (E), and p-S332 CaMKIIδ (F) normalized to total protein; n = 6 Control, 10 HF and LVAD R, 9 HF and LVAD NR; one-way ANOVA with Tukey’s post-hoc test. **(G)** Linear regression analysis of post-LVAD S332 phosphorylation vs. absolute change in LVEF on LVAD therapy; y-axis line at 15 (minimum value to assign positive functional response), x-axis line at 1.5-fold increase in p-S332 vs. non-failing controls; confidence interval = 95%. Data are presented as the mean ± SEM.

### Regulation of CaMKIIδ-B subcellular localization

Since phosphorylation at S332-S334 had previously been shown to prevent nuclear translocation of CaMKIIδ-B in other cell types (*26, 27*), we next sought to determine the role of phosphorylation at these sites in cardiomyocytes. We generated adenovirus vectors expressing GFP-tagged wildtype (B_SSS_), phospho-null (serine to alanine mutations at 332-334 – B_AAA_), and phospho-mimetic (serine to aspartate mutations at 332-334 – B_DDD_) CaMKIIδ-B and transduced neonatal rat ventricular myocytes (NRVMs). As expected, B_DDD_ displayed cytoplasmic localization (**Fig. 4A-B**). However, unlike in non-cardiomyocytes where phospho-null mutations led to complete re-localization of CaMKIIδ-B to the nucleus (*26*), the B_AAA_ version displayed only modest nuclear localization in NRVMs (**Fig. 4A-B**). Notably, when NRVMs were treated with the adrenergic agonist phenylephrine (PE), nuclear localization of B_AAA_ was potently induced, while B_DDD_ remained cytoplasmic (**Fig. 4A-D**). Nuclear translocation was also induced with endothelin-1, but not caffeine or insulin-like growth factor-1 (**Fig. S13**). We therefore hypothesized that both NLS dephosphorylation and CaMKIIδ autoactivation were required for nuclear translocation in cardiomyocytes, as adrenergic and endothelin receptor activation increase cytosolic Ca^2+^ concentration, leading to CaMKIIδ autoactivation by phosphorylation at T287 (*30*–*32*). We assessed markers of CaMKIIδ activation by western blot and found that autophosphorylation at T287 significantly increased with PE (**Fig. 4E-F**), while M281/282 oxidation displayed a modest, but non-significant increase (**Fig. 4E, Fig. S14**).

**Fig. 4.**
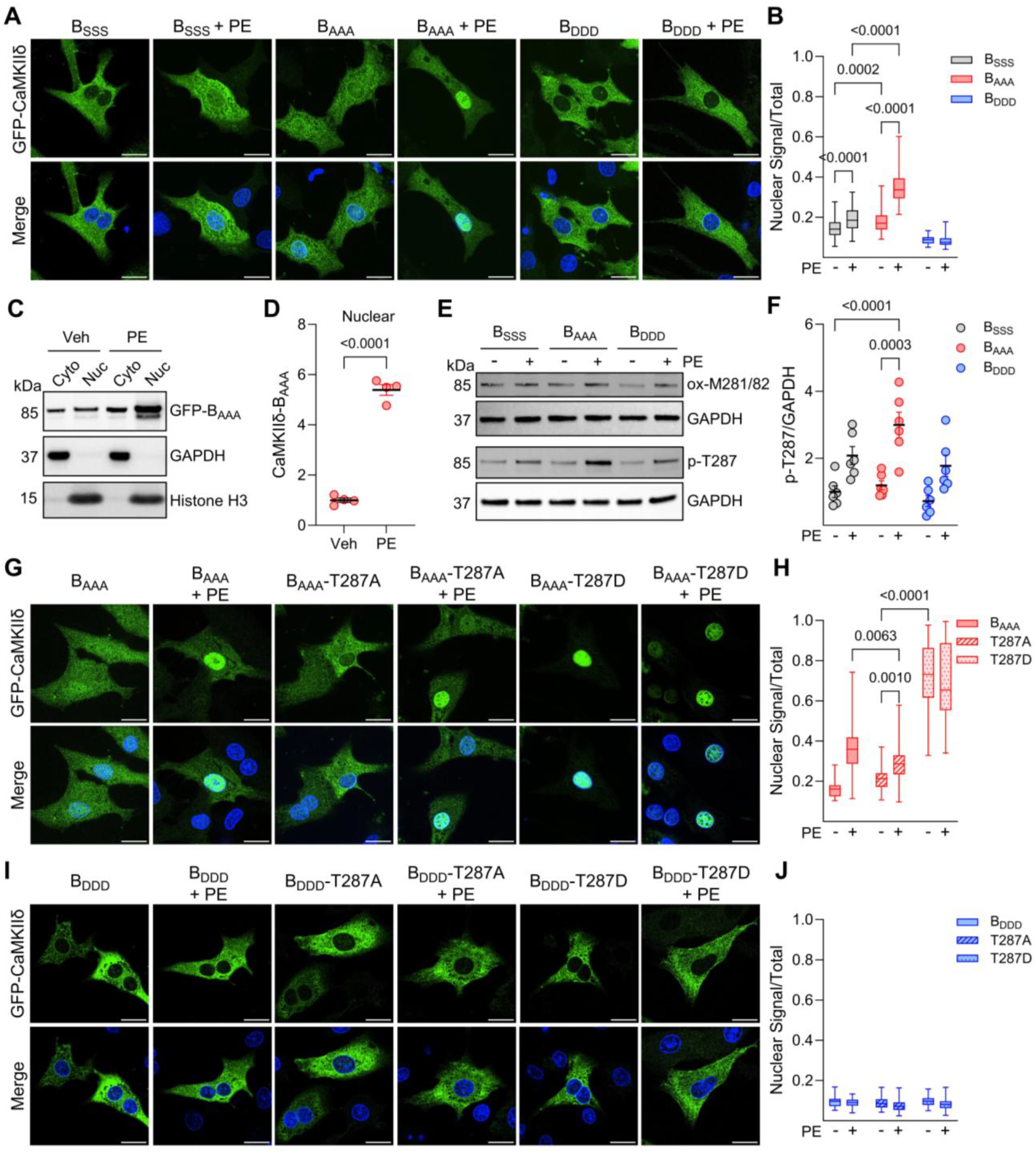
Phosphorylation at S332-S334 prevents autoactivation-dependent CaMKIIδ-B nuclear translocation. **(A)** Representative images for NRVMs 24-hours post-transduction with GFP-CaMKIIδ-B adenoviruses ± PE (20 µM); B_SSS_ = wildtype, B_AAA_ = Serine 332-334 mutated to Alanine (Phospho-null), B_DDD_ = Serine 332-334 mutated to Aspartic Acid (Phospho-mimetic). **(B)** Nuclear GFP normalized to whole-cell GFP; Vehicle: n = 75 B_SSS_, 82 B_AAA_, and 72 B_DDD_. PE: n = 57 B_SSS_ + PE, 72 B_AAA_, 91 B_DDD_ from three biological replicates; two-way ANOVA with Tukey’s post-hoc test. **(C)** Representative western blot for GFP-B_AAA_ in the nuclear and cytosolic fractions of NRVMs treated with vehicle or PE. **(D)** Nuclear GFP-B_AAA_ normalized to Histone H3; n = 4/group; two-tailed t-test. **(E)** Representative western blots for M281/282 oxidized and T287 phosphorylated GFP-CaMKIIδ-B in adenovirus-transduced NRVMs ± PE. **(F)** p-T287 normalized to GAPDH; n = 6/group; two-way ANOVA with Tukey’s post-hoc test. **(G)** Representative images for NRVMs 24-hours post-transduction with B_AAA_ or B_AAA_-T287A/D mutants. **(H)** Nuclear GFP normalized to whole-cell GFP; Vehicle: n = 16 B_AAA_, 82 T287A, and 68 T287D. PE: n = 66 B_AAA_, 91 T287A, and 117 T287D from three biological replicates; two-way ANOVA with Tukey’s post-hoc test. **(I)** Representative images for NRVMs 24-hours post-transduction with B_DDD_ or B_DDD_-T287A/D mutants. **(J)** Nuclear GFP normalized to whole-cell GFP; Vehicle: n = 52 B_DDD_, 40 T287A, and 82 T287D. PE: n = 47 B_DDD_, 67 T287A, and 83 T287D from three biological replicates. Data in B, D, and F are presented as the mean ± SEM. Scale bars for microscopy images = 15 µm.

To test the hypothesis that both T287 autophosphorylation and S332-334 dephosphorylation were required for nuclear translocation, we generated adenovirus expression vectors from the B_AAA_ and B_DDD_ constructs where phospho-null (T287A) or phospho-mimetic (T287D) mutations were made at the autoactivation site. As hypothesized, B_AAA_-T287D displayed near-complete nuclear localization that was PE-independent (**Fig. 4G-H**). Meanwhile, B_DDD_ localized to the cytoplasm and was unaffected by T287A/D modifications or PE treatment (**Fig. 4I-J**), indicating that NLS de-phosphorylation is a pre-requisite for autoactivation-dependent nuclear translocation of CaMKIIδ-B. Interestingly, the B_AAA_-T287A construct did exhibit some nuclear localization with PE treatment, albeit to a lesser extent than B_AAA_ alone (**Fig. 4G-H**). The reason for this is unknown, but we expect that CaMKIIδ-B nuclear translocation, while requiring structural changes in the regulatory domain, may be agnostic to the type of activating mechanism and therefore also responds to M281/282 oxidation (modestly increased with PE, **Fig. 4E**) or S280 O-GlcNAcylation (*33, 34*).

### Cellular signaling and functional consequences of cytoplasmic CaMKIIδ-B

Due to the inherent variability associated with human samples and the stringent statistical cutoffs employed, our phospho-proteomics analysis identified few significant hits, thus limiting insight into potential cellular signaling consequences of increased cytoplasmic CaMKIIδ-B. However, we reasoned that the phosphorylation events mediated (directly or indirectly) by cytoplasmic CaMKIIδ-B would correlate with p-S332-S334 levels. We therefore performed linear regression analysis of the entire phospho-proteomics dataset versus p-S332-S334 CaMKIIδ-B, which identified several previously established CaMKIIδ substrates among the strongest correlated events (**Table S4**). Pathway over-enrichment analyses of phospho-sites with an r^2^ > 0.50 versus p-S332-S334 CaMKIIδ-B identified Rho GTPase signaling, cardiac conduction, and regulation of cardiac hypertrophy as the top affected pathways (**Fig. S15**).

To further explore the cellular signaling changes regulated by cytoplasmic CaMKIIδ-B in a more controlled manner, we performed TMT quantitative phospho-proteomics on NRVMs transduced with B_AAA_ or B_DDD_ and treated with PE or vehicle. This approach identified the baseline phospho-proteome effects of nuclear-competent and cytoplasm-restricted CaMKIIδ-B, as well as autoactivation-dependent effects (**Fig. 5A-E**). At baseline, phosphorylation events that increased in NRVMs transduced with B_AAA_ were enriched with cell surface and cytoskeletal proteins, while the downregulated phosphorylation events included nuclear proteins involved in alternative splicing (**Fig. S16**). With PE treatment, B_AAA_ led to increased phosphorylation of nuclear-localized RNA processing factors (**Fig. 5D, Fig. S17**). PE-dependent phosphorylation events that increased in B_DDD_-transduced cells were enriched with proteins involved in Rho GTPase regulation (**Fig. 5E**), matching the findings in the human phospho-proteome (**Fig. S15**). Additionally, when we mined the human and NRVM phospho-proteomics datasets for shared features, we identified multiple individual peptide examples with similar phosphorylation differences between non-responders/B_DDD_ and responders/B_AAA_ (**Fig. S18**), further supporting that cytoplasmic CaMKIIδ-B remodeled the phospho-proteome of non-responders.

**Fig. 5.**
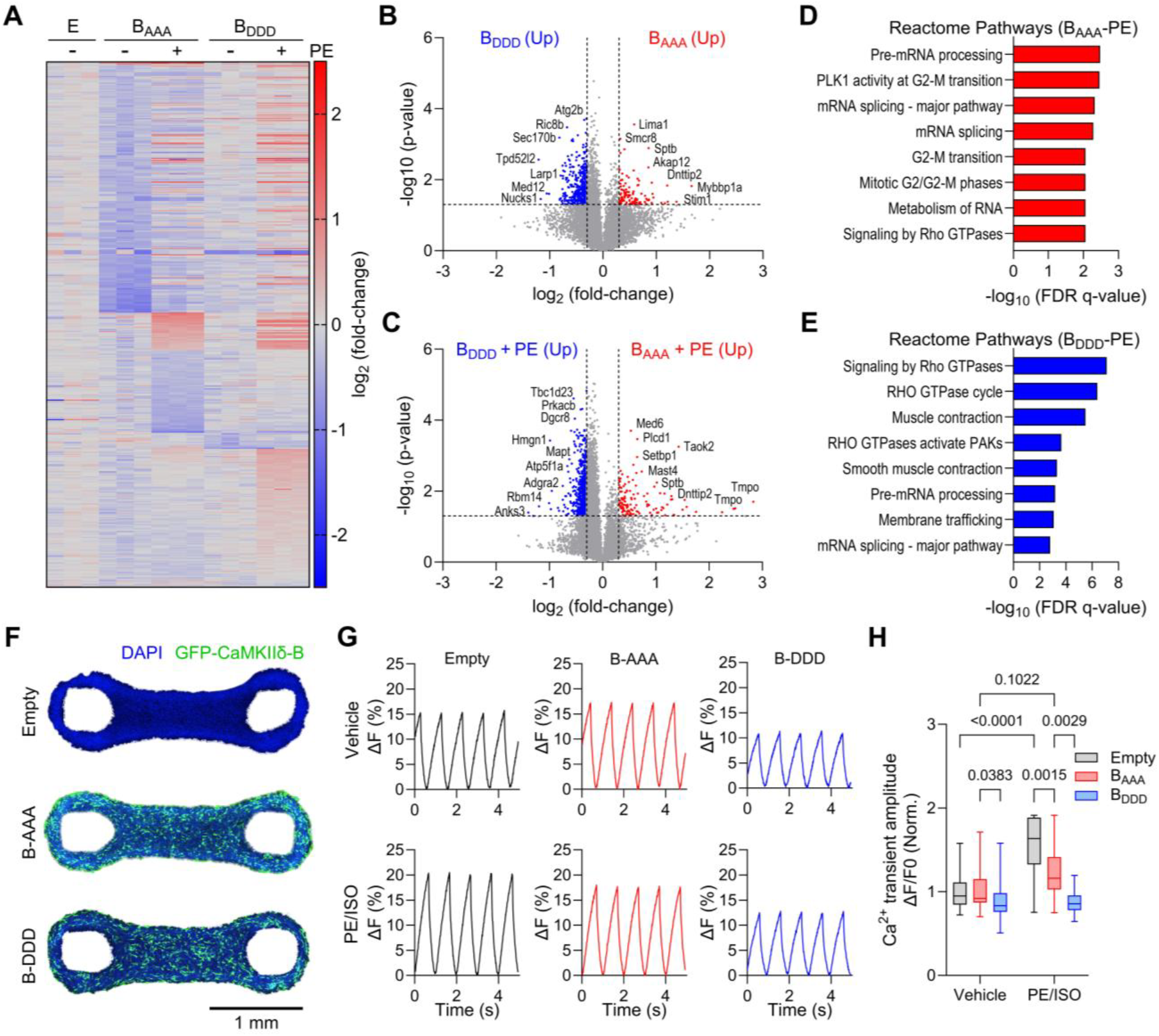
Cytoplasmic CaMKIIδ-B remodels the cardiomyocyte phospho-proteome and impairs contractility. **(A)** Heat map depicting the differentially expressed phospho-peptides in NRVMs transduced with empty vector (E), BAAA, or BDDD ± PE; p-value < 0.01, log_2_ fold-change < -0.3, > 0.3. **(B)** Volcano plot displaying differential phospho-peptide expression in NRVMs transduced with B_AAA_ or B_DDD_. **(C)** Volcano plot displaying differential phospho-peptide expression in NRVMs transduced with B_AAA_ or B_DDD_ and treated with PE. **(D-E)** Reactome Pathway over-enrichment for the differentially expressed phospho-peptides in B_AAA_ + PE (D) versus B_DDD_ + PE (E). **(F)** Representative images of EHTs 24 hours post-transduction with Empty vector, GFP-B_AAA_, or GFP-B_DDD_ adenoviruses; scale bar = 1 mm. **(G)** Representative calcium transient traces for EHTs with and without treatment with the dual α and β adrenergic agonists PE, 50 µM, and isoproterenol (ISO), 2 µM. Traces are plotted as % change in fluorescence (ΔF) over time. **(H)** Mean calcium transient amplitude; Vehicle: n = 41 Empty vector, 43 B_AAA_, 41 B_DDD_ EHTs from three biological replicates. ISO/PE: n = 14 EHTs/group from two biological replicates; two-way ANOVA with Tukey’s post-hoc test.

To test for potential functional consequences of increased cytoplasmic CaMKIIδ-B in human cardiomyocytes, we employed a human induced pluripotent stem cell (iPSC)-derived cardiomyocyte engineered heart tissue (EHT) model (*35*). EHTs were transduced with Empty vector control, B_AAA_, or B_DDD_ adenoviral vectors (**Fig. 5F**) and then Ca^2+^ transient amplitude was measured with and without adrenergic stimulation. EHTs transduced with B_DDD_ had reduced Ca^2+^ transient amplitude compared to B_AAA_ (**Fig. 5G-H**). As expected, adrenergic agonism caused a significant increase in Ca^2+^ transient amplitude in EHTs transduced with the Empty vector control (**Fig. 5G-H**). While the Ca^2+^ transient amplitude in B_AAA_ EHTs did not increase to the same extent as with Empty vector, the response to adrenergic agonism was significantly higher than in B_DDD_ EHTs, which were completely unresponsive to the stimulus (**Fig. 5G-H**).

## Discussion

Restoration of cardiac function in heart failure, while rare and unpredictable, does occur in a small subset of patients treated with current therapies (*8*). Given this evident reversibility of what was long thought of as a static disease state, there is considerable interest in identifying new therapeutic targets for recovery (*8, 36*–*39*). However, a lack of human heart tissue studies paired with molecular mechanistic investigations has resulted in the underlying biology of recovery remaining poorly understood. Previous investigations using multi-omics approaches to identify molecular features of recovery with LVAD identified modest differences in cardiac cell type proportions and gene expression between responders and non-responders (*13, 14*). Our RNA-seq analysis of pre- and post-LVAD samples further corroborated the findings of these previous studies and revealed that the cardiac transcriptome is remarkably similar between these patient sub-groups. However, a deeper analysis at the proteomic and gene isoform levels identified dramatic changes in RNA alternative splicing in heart failure and unique gene isoform expression changes in patients who experienced functional recovery. Among these, alternative splicing and phosphorylation of CaMKIIδ correlated with recovery and we found that this modulation of CaMKIIδ dramatically remodeled the phospho-proteome and impacted cardiomyocyte contractility.

Given the established role of CaMKIIδ in heart disease progression (*24*), the development of therapeutic strategies that inhibit this kinase has been a goal of heart disease research for decades (*22*). However, despite the clear role of CaMKIIδ in driving cardiac pathology, small molecule CaMKII inhibitors have yet to emerge in clinical practice (*22*). Some reasons for this include off-target inhibition of this ubiquitously expressed kinase in other tissues, inhibition of beneficial cardiac CaMKII functions, and failure to discriminate between different CaMKII isoforms (*22*). Recent evidence from pre-clinical studies identified that inhibiting CaMKIIδ constitutive activation by adenine base editing of key residues in the regulatory domain could successfully prevent heart failure development after cardiac injury (*23, 40, 41*). While these findings are exciting, hesitancy around the use of genome editing technology and challenges with adeno-associated virus delivery to the human heart make rapid clinical implementation unlikely, thus warranting the development of additional therapeutic modalities. Our findings indicate that the subcellular localization of CaMKIIδ – specifically mis-localization of the nuclear isoform to the cytoplasm – is a critical driver of the cardiac pathology ascribed to this kinase that has previously been overlooked. Therefore, modulating CaMKIIδ subcellular localization may represent a therapeutic strategy for advanced heart failure. The isoform specificity of such an approach would also mitigate the off-target inhibition concerns posed by current therapies, as the B isoform of CaMKIIδ is primarily expressed in the heart (*42*).

Alternative splicing can generate at least 11 different splice variants from the *CAMK2D* gene (*24*), three of which are highly expressed in cardiomyocytes: δ-B, δ-C, and δ-9 (*24, 43*). In the human heart, the δ-B and δ-9 isoforms comprise ∼90% of total CaMKIIδ (*43*). The δ-C and δ-9 isoforms localize to the cytosol, sarcoplasmic reticulum, and plasma membrane and regulate the activity of proteins involved in Ca^2+^-handling and inflammation (*24, 44*). Overexpression of δ-C or δ-9 in mice induces rapid heart failure onset (*45*–*47*). In contrast, multiple studies have shown that the δ-B isoform, which is the only isoform that includes the NLS encoded by exon 14, has cardioprotective properties. These include prevention of cardiomyocyte apoptosis, inhibition of pro-inflammatory signaling, and induction of mitochondrial Ca^2+^ uptake in conditions of Ca^2+^ overload (*48*–*51*). Overexpression of CaMKIIδ-B does induce hypertrophic cardiac remodeling in mice, but disease onset occurs much later than with overexpression of the cytoplasmic isoforms (*42, 44*). However, considering our findings, it should be noted that simply overexpressing CaMKIIδ-B does not ensure nuclear translocation. Increased CaMKIIδ-B expression in human heart failure, identified herein and in one previous study (*52*), indicates a shift in alternative splicing to a protective isoform. However, our findings show that hyperphosphorylation at the NLS in therapy non-responsive patients prevents CaMKIIδ-B from reaching the nucleus, resulting in increased cytoplasmic CaMKIIδ. Meanwhile, in heart failure patients who experience recovery on LVAD, exon 14 inclusion decreases to healthy control levels as these patients are no longer in the dysfunctional state that triggers this attempted cardioprotective transcriptional response. Our findings suggest that nuclear targeting of CaMKIIδ-B is a molecular strategy to sequester the kinase away from cytoplasmic targets associated with cardiac pathology and, moreover, that restricting this isoform to the cytoplasm prevents recovery from advanced heart failure.

## ACKNOWLEDGMENTS

We thank M. Buvoli, J. Gugel, Y. Tan, and all other members of the Leinwand laboratory for helpful discussions; S. Papasoglu Martin for graphics; W. Wang (University of Washington) for the CaMKIIδ-B p-S332 polyclonal antibody. Funding was provided by the following: United States National Institutes of Health (R01GM029090 to LAL; F32HL170637 to TGM; R01HL16434, R01HL147064, and X01HL139403 to LM and MRGT); American Heart Association (24PRE1195130 to DRH); Shurl and Kay Curci Foundation (Curci Scholars Program Fellowship to CA); Fondation Leducq Transatlantic Network of Excellence (21CVD02 to LAL). Clinical sample data collection and storage was supported by NIH NCATS Colorado CTSA Grant Number UM1 TR004399 (Contents are the authors’ sole responsibility and do not necessarily represent official NIH views). Spinning disc confocal microscopy was performed at the BioFrontiers Institute’s Advanced Light Microscopy Core (RRID: SCR_018302) on a Nikon Ti-E microscope supported by the BioFrontiers Institute and the Howard Hughes Medical Institute. Acrylamide gel and immunoblot imaging were performed with a Cytiva IQ-800 imager in the CU Boulder Biochemistry Shared Instruments Pool (RRID: SCR_018986). The authors acknowledge the BioFrontiers Computing Facility at the University of Colorado Boulder for High Performance Computing and data storage resources supported by BioFrontiers IT. Proteomics analyses were performed at the CU Boulder Proteomics and Mass Spectrometer core facility (RRID: SCR_018992).

## Author Contributions

Conceptualization: TGM, JAB, AVA, PMB, and LAL. Methodology: TGM, DRH, CCE, and APD. Investigation: TGM, DRH, CCE, APD, CA, JCC, SLG, SB, MRB, LM, MRGT, AVA, and PMB. Formal Analysis: TGM, DRH, CCE, and APD. Visualization: TGM and DRH. Funding Acquisition: TGM, LM, MRGT, JAB, LAL. Resources: JCC, SLG, SB, MRB, LM, MRGT, AVA, and PMB. Project Administration: JAB, AVA, PMB, and LAL. Supervision: JAB, LAL. Writing – Original Draft: TGM. Writing – Review and Editing: DRH, CCE, APD, CA, JCC, SLG, SB, MRB, LM, MRGT, AVA, PMB, and LAL.

## Data and Materials Availability

The RNAseq data were deposited at Gene Expression Omnibus (*data will be made public upon acceptance*). Proteomics and phospho-proteomics data were deposited in the PRIDE database within the Proteome Xchange Consortium (*data will be made public upon acceptance*). All other data are available in the main text or associated supplementary materials. Correspondence and requests for materials can be directed to: Dr. Leslie A. Leinwand, BioFrontiers Institute, Jennie Smoly Caruthers Biotechnology Building, D354, 3415 Colorado Ave, Boulder CO, 80303.

## SUPPLEMENTARY MATERIALS

Materials and Methods

Supplemental References 1-13

Figs. S1-S18

Tables S1-S7

Supplemental Data S1-S8

