## Supplemental Information for "Cytoplasmic CaMKIIδ-B prevents myocardial recovery in heart failure"

### Materials and Methods

#### *Clinical Definition of Responders and Non-responders*

Among 41 available paired pre- and post-LVAD patient samples in the University of Colorado Anschutz Medical Center biobank, responder (R) classification (n = 10) was assigned if both of the following criteria were met: 1. The LV ejection fraction (LVEF) increased by an absolute value of > 15 (e.g., from 20% to 35%), and 2. The LV end diastolic dimension (LVEDD) decreased by > 15% following mechanical circulatory support. Non-responder (NR) classification (n = 9) was assigned if both of the following criteria were met: 1. LV ejection fraction increased by an absolute value < 5 or decreased, and 2. LV end diastolic dimension decreased by < 15% or increased following mechanical circulatory support. Patients whose responses did not meet these criteria (n = 22) were considered intermediate responders and were not included in the study.

#### *Human Cardiac Tissue Procurement and Ethical Oversight*

Paired pre-LVAD cores and post-LVAD explanted hearts from 19 patients were obtained with informed consent at the University of Colorado Anschutz Medical Center. All patients receiving LVADs were in New York Heart Association (NYHA) Class IV heart failure (end-stage) at the time of device placement. In all cases, LVAD therapy was used as bridge to transplant. Non-failing control LV tissue (n = 6) was obtained from age- and sex-matched donor hearts without a history of coronary artery disease that were deemed unsuitable for transplant due to size or other incompatibility. Following cardiac explant in the operating room, the samples were flash frozen in liquid nitrogen and then stored at -80°C prior to experimentation. De-identified patient clinical data was stored in a secure REDCap database. Heart failure patient samples were sequenced to identify variants in cardiomyopathy-associated genes, which are reported in **Table S5**. The collection, storage, and experimentation on these samples were approved by the Institutional Review Board of the University of Colorado Anschutz Medical Center.

#### *Animal Studies and Ethical Oversight*

Animal experiments were approved by the Institutional Animal Care and Use Committee (IACUC) at the University of Colorado Boulder. The University of Colorado Boulder is an AAALAC-accredited institution.

#### *Neonatal Rat Ventricular Myocyte Isolation*

NRVMs were isolated from 1–2-day-old Sprague-Dawley rat pups as previously described (*1*). In brief, rat pups were dipped in ethanol, decapitated, and their hearts rapidly removed and placed in calcium and bicarbonate-free HEPES-buffered Hanks' Solution (CBFHH). The atria were then removed and the ventricles from all rats pooled in a beaker containing 10 mL warm CBFHH, Trypsin (3 mg/mL), and 100 µl DNase. The solution was placed on a stir plate and incubated for 15 minutes. Cardiomyocytes were dissociated by multiple sequential incubations in this solution paired with mechanical dissociation by pipetting up and down. When the cells were fully dissociated, they were suspended in recovery media (MEM, 1X PB12, 1X HEPES, 5% calf serum) and pre-plated for 90 minutes to remove adherent cells (e.g., fibroblasts, endothelial cells). The suspended cells (cardiomyocytes) were then plated on gelatin-coated tissue culture plates and incubated at 37°C, 1.5% $CO_2$. The day after isolation, the media was switched to experimental medium (MEM, 1X PB12, 1X HEPES, 1X BrdU, Pen-Strep: 500 mL MEM, 1 mL

PB12 stock, 10 mL 1M HEPES pH 7.3, 1 mL BrdU, 2 mL Pen-Strep stock) and experiments were started two days post-plating.

***RNA Extraction, cDNA Synthesis, and qPCR Analysis***

Frozen left ventricular tissue (~40 mg) was added to 1 mL of Trizol, homogenized with a mechanical homogenizer, and incubated at room temperature for 5-10 minutes. Following this incubation, the homogenate was transferred to a 1.5 mL Eppendorf tube, chloroform added at 1:5 vol/vol, the tubes shaken vigorously for ~15 seconds, and then incubated at room temperature for 15 minutes. The samples were then centrifuged for 15 minutes at 12,000 x g, 4°C and the upper aqueous phase collected and transferred to a clean tube. An equal volume of isopropanol (500 µl) was added, the tubes briefly vortexed, and then incubated at -20°C for 15 minutes followed by centrifugation at 12,000 x g, 4°C for 8 minutes to pellet the RNA. The supernatant was discarded, and RNA pellet washed twice with ice-cold 75% ethanol. After removal of the final ethanol wash, the RNA was dried for 5 minutes by uncapping the tubes and then reconstituted in Milli-Q $H_2O$. RNA concentrations were determined by Nanodrop. 500 ng of RNA was used to synthesize complementary DNA (cDNA) with SuperScript III Reverse Transcriptase (Invitrogen) according to the manufacturer's protocol. The cDNA was then diluted to 2.5 ng/µl and 5 ng total cDNA used for quantitative real-time PCR with the primers listed in **Table S6**. Gene expression analysis was performed using the ΔΔ-CT method with *PGK1*, chosen based on stable high expression across groups in RNA-seq analysis, as the housekeeping gene.

***RT-PCR and DNA Acrylamide Gel Visualization***

PCR was performed using Taq PCR Master Mix (Qiagen). Reactions were 25 µl total volume, including 30 ng of cDNA template. 30 cycles of PCR with the following conditions were conducted: Denaturation – 95°C, 30 seconds; Annealing – 5°C below primer $T_m$, 30 seconds; Extension – 72°C, 1 minute. The primer sequences are listed in **Table S7.** Results were visualized by adding 5 µl of Hi-Density TBE Sample Buffer (Novex) and separating PCR products via electrophoresis on a 10% TBE acrylamide gel at 145V for 1.5 hours. The gel was incubated in 1X ethidium bromide in milliQ-$H_2O$ for 5 minutes and then imaged using the UV setting on an IQ-800 imager (Cytiva).

***Bulk RNA Sequencing and Alternative Splicing Analysis***

RNA was extracted from frozen left ventricular tissue as described above. mRNA was purified by poly-A enrichment, cDNA libraries synthesized, and bulk RNA-seq performed using Illumina NovaSeq with 150-bp paired-end reads (Novogene Corporation, Sacramento, CA). The minimum sequencing depth was 50 million read pairs. Differential gene expression analysis was performed using DESeq2 version 1.46.0 (*2*). For analysis between non-failing controls, HF, and LVAD groups, the dataset was prefiltered to remove genes that had less than 5 samples (size of the smallest group: controls) with 10 or more counts. The apeglm shrinkage estimator (*3*) was used to produce shrunken log fold changes for visualization. Genes were considered significantly differentially expressed if they displayed >1.0, <-1.0 shrunken $\log_2$ fold-change and an FDR-adjusted p-value < 0.05. For comparison of responders and non-responders at pre- and post-LVAD timepoints, genes were considered significantly differentially expressed if they displayed FDR-adjusted p-value < 0.05. To contrast the responses to LVAD treatments between responders and non-responders, a model matrix with the following design was constructed: response + response:patient.n + response:timepoint (see DESeq2 vignette for further details). The dds object was prefiltered to remove genes that had less than 9 samples (size of the smallest group: non-responders) with 10 or more counts. DESeq2 was used to determine differences in the LVAD treatment effect between responders and non-responders, considering the paired nature of the samples. Pathway over-enrichment analysis on the significantly differentially expressed genes was performed using Enrichr (*4*). Pathways with FDR q-values < 0.05 were considered significantly enriched. Gene Set Enrichment Analysis was performed with GSEA (*5*, *6*) version 4.3.3 on pre-ranked gene lists ranked by the negative log of the p-value multiplied by the sign of the fold-change. Majiq HET and Voila version 2.4.dev3+g85d0781 (*7*) were used to analyze the short-read data for alternative splicing differences. Modules were extracted via the Voila Modulize command. Local splicing variations (LSVs) were considered changed if they had a dpsi change greater than 0.15 between conditions and a Wilcoxon p-value < 0.05. For defining modules as changing, default values were used except fore\ the following flags: --changing-between-group-dpsi 0.15 --changing-between-group-dpsi-secondary 0.0. DIGGER (*8*, *9*) (https://exbio.wzw.tum.de/digger/nease_analysis) was used run NEASE analysis on cassette exons that underwent differential splicing (defined as a cassette exon event from Voila Modulize with a changing junction, dpsi > 0.15 and Wilcoxon p-value < 0.05). Majiq HET output was manually converted to the standard input format for DIGGER NEASE analysis.

***Protein Extraction for Proteomics Studies***

*Tissue.* Post-LVAD left ventricular tissue (n = 9 responders, 9 non-responders) was pulverized while frozen, solubilized in a 5% SDS, 0.75% sodium deoxycholate, 50 mM Tris pH 8 buffer containing 1X protease and phosphatase inhibitors (Halt Protease and Phosphatase Inhibitor Cocktail, Thermo) and immediately heated for 10 minutes 95°C. The samples were then centrifuged at 12,000 x g for 10 minutes to remove any remaining insoluble material and a BCA assay was performed on the soluble portion to determine protein concentration. *Cells.* NRVMs on 10-cm tissue culture plates (5.5 million cells per plate) were washed twice with PBS and then rapidly frozen at -80°C before thawing and lysing in 5% SDS, 0.75% sodium deoxycholate, 50 mM Tris pH 8 buffer containing 1X protease and phosphatase inhibitors by scraping. Lysates from two 10-cm plates were pooled to ensure >1 mg of total protein and make up one biological replicate. Collected lysates were then immediately heated for 10 minutes 95°C. The samples were then centrifuged at 12,000 x g for 10 minutes to remove any remaining insoluble material and a BCA assay was performed on the soluble portion to determine protein concentration.

***Tandem Mass Tag (TMT) Quantitative Proteomics and Phospho-proteomics***

Human heart tissue protein lysates (1300-1800 ug) and NRVM protein lysates (350-550 µg) were reduced and alkylated with the addition of 10 mM tris(2-carboxyethylphosphine) (TCEP), 40 mM 2-chloroacetamide, incubated at 70°C for 10 minutes and then shaking at 2000 rpm at 37°C for 30 minutes. Lysates were digested using the SP3 method (*10*). Briefly, 1000 µg carboxylate-functionalized speedbeads (Cytiva Life Sciences) were added to protein lysates. Acetonitrile was added to 80% (v/v) to bind proteins to the beads, then washed twice with 80% (v/v) ethanol and twice with 100% acetonitrile. Proteins were digested in 50 mM Tris-HCl buffer, pH 8.5, with Lys-C/Trypsin (Promega) and incubated at 37˚C overnight. Tryptic peptides were desalted using HLB Oasis cartridges (Waters) according to the manufacturer's instructions and dried in a speedvac vacuum centrifuge. Phosphopeptides were serially enriched using the High-Select TiO2 phosphoenrichment kit (Thermo Scientific) followed by the High-Select Fe-NTA phosphoenrichment kit (Thermo Scientific). Briefly, samples were dissolved in

Binding/Equilibration buffer, loaded on the TiO2 column, washed and eluted. The TiO2 elutions and unretained fraction were dried immediately using a speedvac vacuum centrifuge. The TiO2 unretained fraction was then dissolved in Fe-NTA Binding/Wash Buffer, loaded on the Fe-NTA column, washed and eluted. The TiO2 and Fe-NTA elutions were combined and cleaned up then labeled with TMT-Pro reagents (Thermo Scientific) along with proteome samples from human hearts according to the manufacturer's instructions. Multiplexed samples were immediately cleaned up with HLB Oasis cartridge and dried in a speedvac vacuum centrifuge. To reduce sample complexity, multiplexed peptides were fractionated using a high pH reversed-phase C18 UPLC with a 0.5 mm X 150 mm custom packed rpC18 1.9 µm 120Å (Dr. Maisch Gmbh) column with mobile phases 0.1% (v/v) aqueous ammonia, pH10 in water and acetonitrile (ACN). Peptides were gradient eluted at 20 µL/minute from 2 to 50% ACN in 50 minutes (proteome) or 2 to 20% ACN in 50 minutes (phosphoproteome) concatenating for a total of 12 fractions using a Waters M-class UPLC (Waters). Peptide fractions were then lyophilized in a speedvac vacuum centrifuge and stored at -20°C until analysis. Fractionated peptides were suspended in 3% (v/v) ACN, 0.1% (v/v) trifluoroacetic acid (TFA) and directly injected onto a reversed-phase C18 1.7 µm, 130 Å, 75 mm X 250 mm M-class column (Waters), using either an Ultimate 3000 nanoUPLC coupled to a Q Exactive HF-X (human heart tissues) or a Vanquish Neo nanoUPLC coupled to an Orbitrap Exploris 480 (NRVMs) (Thermos Scientific). Multiplexed peptides were eluted into the mass spectrometers at 300 nL/minute with a gradient from 4% to 16% ACN in 120 minutes then to 40% ACN in 5 minutes (human hearts) or 4% to 20% ACN in 120 minutes then to 40% ACN in 5 minutes (NRVMs). Precursor mass spectra (MS1) were acquired at a resolution of 120,000 from 350 to 1500 m/z with an automatic gain control (AGC) target of 3E6 and a maximum injection time of 50 milliseconds (human hearts) or Standard AGC Target and Auto Maximum injection time (NRVMs). Precursor peptide ion isolation width for MS2 fragment scans was 0.7 m/z, and the top 15 most intense ions were sequenced (human hearts) or a cycle time of 3 seconds (NRVMs). MS2 spectra were either acquired at a resolution of 45,000 with higher energy collision dissociation (HCD) at 30% normalized collision energy (human hearts) or at 30,000 resolution using TurboTMT with HCD at 32%. An AGC target of 1E5 and 120 milliseconds maximum injection time (human hearts) or 200% Normalized AGC Target and Auto maximum injection time (NRVMs) were used. Dynamic exclusion was set for 30 seconds. Rawfiles were searched against either the Human database (UP000005640) using MaxQuant v.2.0.3.0 or the Rattus norvegicus database (UP000002494) using MaxQuant v.2.6.3.0. Cysteine carbamidomethylation was considered a fixed modification, while methionine oxidation, protein N-terminal acetylation and phosphorylation of serine, threonine, or tyrosine were searched as variable modifications. All peptide and protein identifications were thresholded at a 1% false discovery rate (FDR). TMT reporter ion intensities were Cyclic loess normalized and log2 fold changes and p-values were calculated with limma (Bioconductor.com) using an R-script. Pathway over-enrichment analysis on the significantly differentially expressed proteins was performed using Enrichr (*4*). Pathways with FDR q-values < 0.05 were considered significantly enriched.

***Subcellular Fractionation***

NRVMs on 60-mm tissue culture plates were washed twice with PBS and then 300 µl of buffer (10 mM Tris-HCl pH 7.4, 10 mM NaCl, 3 mM $MgCl_2$, 0.3% NP-40, 1X protease/phosphatase inhibitors) added and cells collected by scraping. The cells in buffer were added to Eppendorf tubes and incubated on ice for 3 minutes. Nuclei were then pelleted by centrifugation at 800 x g for 5 minutes. The supernatant was collected as the crude cytosolic fraction, which was then clarified by centrifugation at 12,000 x g for 10 minutes. The nuclear pellet was washed twice in buffer without NP-40 and then resuspended in 5% SDS, 0.75% deoxycholate buffer and heated at 95°C for 10 minutes to lyse.

***Protein Extraction for Western Blot Analyses***

*Tissue:* Left ventricular tissue was pulverized while frozen, added to lysis buffer (5% SDS, 0.75% sodium deoxycholate, 50 mM Tris pH 8) containing 1X protease and phosphatase inhibitors, and then immediately heated for 10 minutes 95°C. Samples were then centrifuged at 12,000 x g for 10 minutes, 4°C and the supernatant collected. *Cells:* NRVMs on 6-well tissue culture plates were washed twice with PBS and then plates were immediately frozen at -80°C. Plates were brought to room temperature and then 150 µl lysis buffer (5% SDS, 0.75% deoxycholate, as above) containing 1X protease and phosphatase inhibitors added to the wells. Cells were lysed with a cell scraper and then the collected material immediately heat at 95°C for 10 minutes. Samples were then centrifuged at 12,000 x g for 10 minutes, 4°C and the supernatant collected.

***Western Blot***

Protein concentration was determined by BCA assay and then 20 µg of protein was added to equal volume loading buffer (SDS Tris Glycine buffer + Bolt Reducing Agent) and heated at 95°C for 8 minutes. Proteins were loaded onto 4-12% Bis-Tris gradient gels (Invitrogen) for separation by electrophoresis and then transferred onto nitrocellulose membrane. To assess loading and quantify total protein prior to blocking, membranes were incubated in Revert Total Protein stain (LICOR) for 5 minutes followed by image acquisition on an IQ-800 imager (Cytiva) using the IR-short setting with automatic exposure. Membranes were blocked in 5% milk TBS-T for 1 hour at room temperature and then primary antibodies were added in 5% BSA TBS-T and incubated overnight at 4°C with agitation. Primary antibodies and dilutions: CaMKIIδ (Thermo Fisher, PA5-22168, 1:1000), Phospho-CaMKIIδ T287 (Cell Signaling, 12716, 1:1000), Phospho-CaMKIIδ S332 (a gift from Dr. Wang Wang, University of Washington, 1:100), Histone H3 (Cell Signaling, 4499, 1:1000), GAPDH (Cell Signaling, 2118, 1:1000). Following overnight incubation, primary antibodies were removed, the membranes washed three times for 5 minutes each with TBS-T, and secondary antibodies (IRDye CW800 goat anti-rabbit, LICOR) added in 5% Milk TBS-T at 1:4000 dilution and incubated for 1 hour at room temperature with agitation. Secondary antibodies were then removed, the membranes washed three times for 5 minutes each with TBS. Blots were imaged on an IQ-800 imager using the IR-long setting and quantification of signal intensity was performed using Fiji.

***Cloning and Adenovirus Production***

The Human CAMK2D Gene ORF cDNA clone expression plasmid (Catalog #: HG29766-NF), encoding the CAMK2D (minus exons 14-16 and 21-22), was purchased from Sino Biological. Molecular cloning with Gibson Assembly was used to add a synthetic double-stranded oligo encoding exons 21 and 22 to the C-terminus and an N-terminal EGFP. The resulting EGFP-CAMK2D-full-length ORF was assembled in the pShuttle-CMV shuttle vector by Gibson Assembly. Cloning was confirmed by sequencing. The pShuttle-CMV-EGFP-CAMK2D plasmid was then used as a PCR template with Gibson Assembly to generate the wild-type CAMK2D-B isoform (addition of 33 base pairs encoding exon 14), phospho-null CAMK2D-$B_{AAA}$ (AAA –

serine to alanine mutations at amino acids 332, 333, and 334), phospho-mimetic CAMK2D-$B_{DDD}$ (DDD – serine to aspartate mutations at amino acids 332, 333, and 334). The $B_{AAA}$ and $B_{DDD}$ constructs were then used as PCR templates with Gibson Assembly to generate the T287A/D mutants. pShuttle-CMV empty vector was prepared as a negative control. All cloning was confirmed by whole plasmid sequencing from Plasmidsaurus. pShuttle constructs were then subcloned into the pAdEasy backbone and used to transfect HEK293 cells. Ten days after transfection, the virus was passaged onto new plates and then amplified in subsequent larger scale infections. Purified viruses for each were obtained from 72 two-day post-infection 10-cm plates by overlaying concentrated freeze-thawed cell lysates on CsCl gradients and ultracentrifuging. Adenoviral particles were extracted from Beckman centrifuge tubes by side-puncture with an 18-gauge needle and stored in 1:4 ratio in Falck-Pedersen viral storage buffer at -20°C. A multiplicity of infection (MOI) of 20 was used for *in vitro* experiments with these viruses in NRVMs and iPSC-CM EHTs.

***Adenovirus Transduction Experiments in NRVMs***
Two days after their isolation, NRVMs were transduced with adenoviral constructs expressing empty vector or EGFP-tagged $B_{SSS}$, $B_{AAA}$, $B_{DDD}$, $B_{AAA}$-T287A, $B_{AAA}$-T287D, $B_{DDD}$-T287A, or $B_{DDD}$-T287D at 20 viral particles per cell (20 MOI). Phenylephrine (PE) was added to the media at this timepoint at a final concentration of 20 µM. 24 hours after viral transduction/PE treatment, protein lysates were collected for western blot/phospho-proteomic analyses or cells were fixed for immunofluorescence microscopy analysis of GFP-CaMKIIδ-B localization. For western blot experiments, NRVMs were plated on 6-well dishes at 500,000 cells per well. For immunofluorescence microscopy experiments, NRVMs were plated on 24-well glass-bottomed dishes at 100,000 cells per well. For phospho-proteomics experiments, 5.5 million cells were plated on 10-cm tissue culture plates.

***Immunofluorescence Microscopy***
Cells or EHTs were washed twice with PBS and then fixed in ice-cold 100% methanol for 1 minute, followed by 4% paraformaldehyde for 5 minutes. Permeabilization was performed using 0.5% triton in PBS for 20 minutes at room temperature. Antigen retrieval was then conducted by incubation in 100 mM glycine, pH 3.5, for 30 minutes at room temperature. The wells were washed three times with PBS and then incubated in blocking buffer (5% BSA in PBS – 0.25 µm filtered) for 1 hour at room temperature. Mouse α-actinin primary antibody (Clone EA-53, Thermo) was added in fresh blocking buffer at 1:300 dilution and incubated overnight at 4°C. The following day, the wells were washed three times with PBS and Alexa-Fluor goat anti-mouse 555 secondary antibody (4409, Cell Signaling) added in fresh blocking buffer at 1:1000 dilution. The plate was then incubated for 1 hour at room temperature protected from the light. Following this incubation, the wells were washed twice with PBS and then DAPI (D3571, Thermo) added in PBS and incubated for 15 minutes at room temperature. Two additional washes were performed and then 0.5 mL PBS added per well prior to imaging on a Nikon Ti-E spinning disc confocal microscope. Images were acquired using constant laser intensity and photomultiplier gain settings. Intensity of nuclear and whole-cell GFP signal was measured using Fiji (Image J). Nuclear GFP-CaMKIIδ-B intensity was normalized to total GFP signal from the same cell for quantification.

***iPSC-derived Cardiomyocyte Differentiation and Culture***

Human induced pluripotent stem cells (WTC11) were grown in E8 medium on Matrigel-coated plates and split twice before media switch to RPMI 1640 for induction of cardiomyocyte differentiation. Mesoderm differentiation was induced in RPMI containing B27 minus insulin, 50 µg/mL ascorbic acid, and 7 µM CHIR-99021 (GSK3 inhibitor) for 48 hours. The cells were allowed to recover in RPMI/B27 minus insulin/ascorbic acid medium for one day and then cardiac mesoderm specification was induced using 5 µM IWR1 (Wnt inhibitor) in RPMI/B27 minus insulin/ascorbic acid medium for 48 hours. The cells were allowed to recover for one day, as above, then media was changed to RPMI with B27 plus insulin. Beating cells were observed on day 7-8 after GSK3 inhibition. The cultures were maintained in RPMI/B27 plus insulin medium until day 12 when they were collected and seeded into engineered heart tissue (EHT) molds.

***Human Engineered Heart Tissue (EHT) Generation***

Hydrogel micropillar molds were 3D printed using a custom poly(ethylene glycol) diacrylate resin on a Lumen Alpha DLP printer (Volumetric Inc., USA) with predefined light exposure settings (7.5 s per layer, 100 µm layer thickness, 20 mW $cm^{-2}$) as described previously (*11*, *12*). Following 3D printing, hydrogel micropillar molds were washed in DPBS (with multiple rinses over 3-5 days to remove unreacted reagents), sterilized with immersion in 70% ethanol for 30 minutes followed by UV germicidal lamp for another 30 minutes, and stored in sterile DPBS (1X, Gibco) at 4°C until use. The stiffness of the micropillar was determined to be 2.096 µN $µm^{-1}$ via static stress simulation of micropillar bending using Fusion 360 (Autodesk, USA). To form EHTs, hydrogel micropillar molds were seeded with iPSC-derived cardiomyocytes (cell age: day 12, density: 10 million cells $mL^{-1}$). Briefly, acid-solubilized type I bovine telocollagen (Advanced BioMatrix, USA) was mixed with prechilled, sterile DPBS (10X), distilled water, Matrigel® (basement membrane matrix growth factor reduced, 10% *v/v*, Corning), and neutralized with NaOH (0.1 N) to achieve a final collagen concentration of 1 mg $mL^{-1}$. Cells were added to the mixture, the resultant precursor (6 µL) was transferred into micropillar wells under ice-cold conditions, and the molds were incubated (37°C, 5% $CO_2$ incubator) for 15 min to enable collagen gelation. After incubation, the molds containing cell-laden gels were immersed in recovery medium comprising RPMI 1640 (Gibco) supplemented with B-27 (1X, Thermo Scientific), FBS (20% v/v), and penicillin–streptomycin (1% v/v). Two days after seeding, the recovery medium was replaced with growth medium consisting of RPMI 1640 with B-27 (1X) and penicillin–streptomycin (1% v/v) with regular media changes every other day. After growth phase (day 10 after seeding), the media was switched to maturation media consisting of low glucose RPMI supplemented with palmitic acid (PA, 50 $\mu$M), oleic acid (OA, 100 $\mu$M), galactose (10 mM), B27 (1X), and penicillin–streptomycin (1% v/v). EHTs were cultured in maturation media until day 28 (i.e., cell age: day 40) before initiating treatment.

***Calcium/Contractility Analysis in Engineered Heart Tissues***

On day 28 after seeding (i.e., cell age: day 40), EHTs were transduced with adenoviruses at 20 MOI. One day later, a subset of EHTs were treated with a combination of the adrenergic agonists phenylephrine (PE, 50 µM) and isoproterenol (ISO, 2 µM). 48 hours after viral transduction, and 24 hours after agonist addition, EHTs were washed with sterile DPBS (1X) and incubated (37°C, 5% $CO_2$ incubator, 90 min) with intracellular calcium indicator (Cal 520, AAT Bioquest, 4 µM) in sterile Tyrode's salt solution (Sigma) supplemented with Pluronic F-127 (0.04 wt.%, Sigma) to enhance cellular uptake as described previously (*12*, *13*). Following dye loading, EHTs were washed and incubated in fresh Tyrode's solution to remove the staining solution. Live cell videos of EHT contractility were recorded on a Nikon Ti2 Eclipse with AXR laser scanning confocal microscope (2X objective, 105 Hz frame rate, 37°C with 5% $CO_2$) under electrical stimulation (1 Hz, 10 V $cm^{-1}$, carbon electrodes with Myopacer setup, IonOptix). All EHTs were first allowed to reach equilibrium by subjecting to stimulation frequency for 5 min before recording any measurements. Calcium transient traces as recorded by the temporal change in fluorescence intensity upon stimulation were used for all measurements. Analysis of parameters (i.e., mean calcium amplitude) was performed using an automated software (Beatprofiler) as described previously (*14*).

***Statistical Analysis and Data Presentation***

Comparisons of two groups were performed by paired or unpaired two-tailed t-test, as described in the figure legends. Paired analyses were performed for pre- versus post-LVAD comparisons and unpaired analyses for pre- versus pre-LVAD and post- versus post-LVAD comparisons pertaining to the responder and non-responder groups. Comparisons of more than two groups were performed by one- or two-way ANOVA, as described in the figure legends. Comparisons of categorical data between responders and non-responders pre- and post-LVAD were performed using the Chi-squared test. All data analyses and graphical representations were performed using GraphPad Prism version 10. The data throughout the manuscript are presented as the mean ± standard error, unless otherwise noted. For the RNA-seq and proteomics data and the corresponding gene ontology enrichment analyses, an adjusted p-value of $< 0.05$ was considered statistically significant. For all other data analyses, a p-value of $< 0.05$ was considered statistically significant.

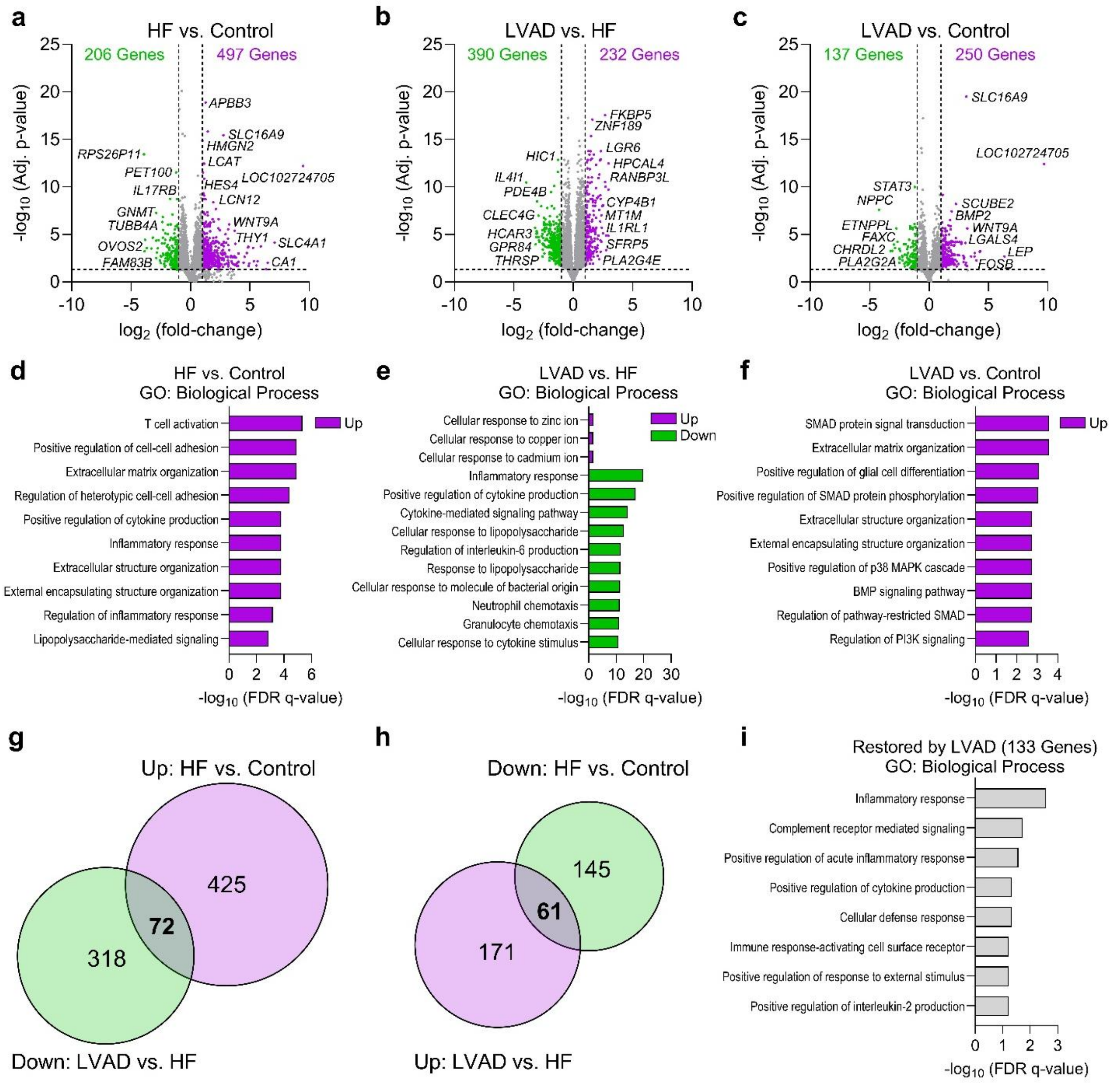
a
HF vs. Control
206 Genes
497 Genes
-log10 (Adj. p-value)
log2 (fold-change)
APBB3
SLC16A9
HMGN2
LCAT
LOC102724705
HES4
LCN12
WNT9A
THY1
SLC4A1
CA1
RPS26P11
PET100
IL17RB
GNMT
TUBB4A
OVOS2
FAM83B
b
LVAD vs. HF
390 Genes
232 Genes
FKBP5
ZNF189
LGR6
HPCAL4
RANBP3L
CYP4B1
MT1M
IL1RL1
SFRP5
PLA2G4E
HIC1
IL4I1
PDE4B
CLEC4G
HCAR3
GPR84
THRSP
c
LVAD vs. Control
137 Genes
250 Genes
SLC16A9
LOC102724705
STAT3
NPPC
ETNPPL
FAXC
CHRDL2
PLA2G2A
SCUBE2
BMP2
WNT9A
LGALS4
LEP
FOSB
d
HF vs. Control
GO: Biological Process
Up
T cell activation
Positive regulation of cell-cell adhesion
Extracellular matrix organization
Regulation of heterotypic cell-cell adhesion
Positive regulation of cytokine production
Inflammatory response
Extracellular structure organization
External encapsulating structure organization
Regulation of inflammatory response
Lipopolysaccharide-mediated signaling
-log10 (FDR q-value)
e
LVAD vs. HF
GO: Biological Process
Up
Down
Cellular response to zinc ion
Cellular response to copper ion
Cellular response to cadmium ion
Inflammatory response
Positive regulation of cytokine production
Cytokine-mediated signaling pathway
Cellular response to lipopolysaccharide
Regulation of interleukin-6 production
Response to lipopolysaccharide
Cellular response to molecule of bacterial origin
Neutrophil chemotaxis
Granulocyte chemotaxis
Cellular response to cytokine stimulus
-log10 (FDR q-value)
f
LVAD vs. Control
GO: Biological Process
Up
SMAD protein signal transduction
Extracellular matrix organization
Positive regulation of glial cell differentiation
Positive regulation of SMAD protein phosphorylation
Extracellular structure organization
External encapsulating structure organization
Positive regulation of p38 MAPK cascade
BMP signaling pathway
Regulation of pathway-restricted SMAD
Regulation of PI3K signaling
-log10 (FDR q-value)
g
Up: HF vs. Control
425
72
318
Down: LVAD vs. HF
h
Down: HF vs. Control
145
61
171
Up: LVAD vs. HF
i
Restored by LVAD (133 Genes)
GO: Biological Process
Inflammatory response
Complement receptor mediated signaling
Positive regulation of acute inflammatory response
Positive regulation of cytokine production
Cellular defense response
Immune response-activating cell surface receptor
Positive regulation of response to external stimulus
Positive regulation of interleukin-2 production
-log10 (FDR q-value)

**Fig. S1. Mechanical circulatory support partially reverses transcriptomic features of heart failure.** a-c. Volcano plots depicting differentially expressed genes identified by RNA-seq between heart failure (HF) and non-failing controls (a), LVAD and HF (b), and LVAD and controls (c). d-f. Significantly enriched GO: Biological Process Pathways for the upregulated and downregulated genes between the three comparisons; enriched pathways only reached significance for the upregulated genes in the HF vs. Control and LVAD vs. Control comparisons. g. Venn Diagrams depicting significantly differentially expressed genes that increased with HF and were reversed with LVAD. h. Venn Diagrams depicting significantly differentially expressed genes that decreased with HF and were reversed with LVAD. i. GO: Biological Process pathway enrichment for genes whose expression was restored by LVAD to Control expression levels. For all RNA-seq, n = 5 Control, n = 19 HF, n = 19 LVAD.

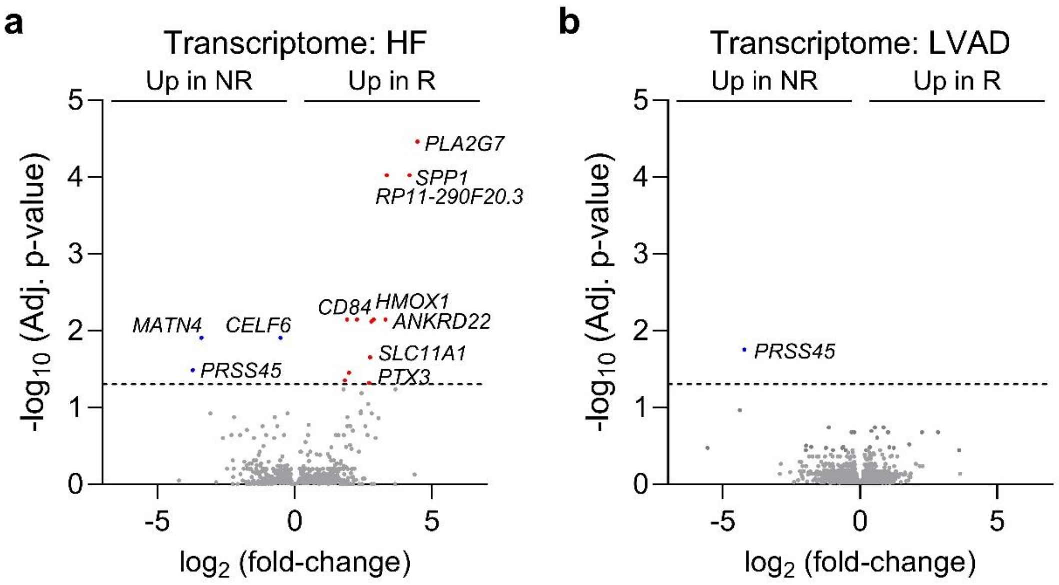


**Fig. S2. Responders and non-responders display similar gene expression profiles pre- and post-LVAD. a-b.** Volcano plots depicting differentially expressed genes identified by RNA-seq between responders (R) and non-responders (NR) in heart failure (a) and post-LVAD (b). An FDR adjusted p-value cutoff of 0.05 was used; n = 9 NR, 10 R. X-axis represents the shrunken $\log_2$ fold-change using the apeglm algorithm.

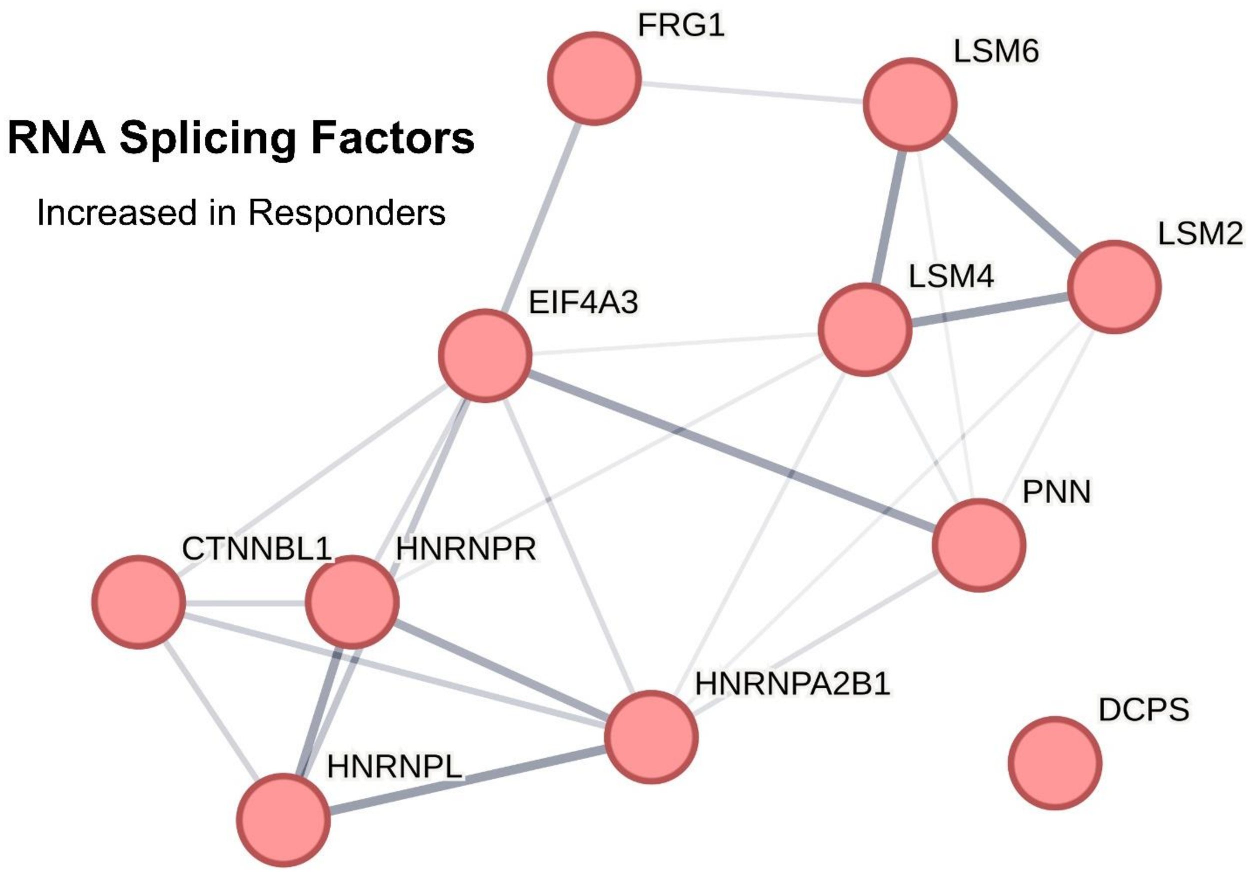


**Fig. S3. Interactome of RNA splicing factors increased in LVAD responders.** STRING analysis of the 11 RNA splicing factors increased in post-LVAD responders identified by quantitative proteomics.

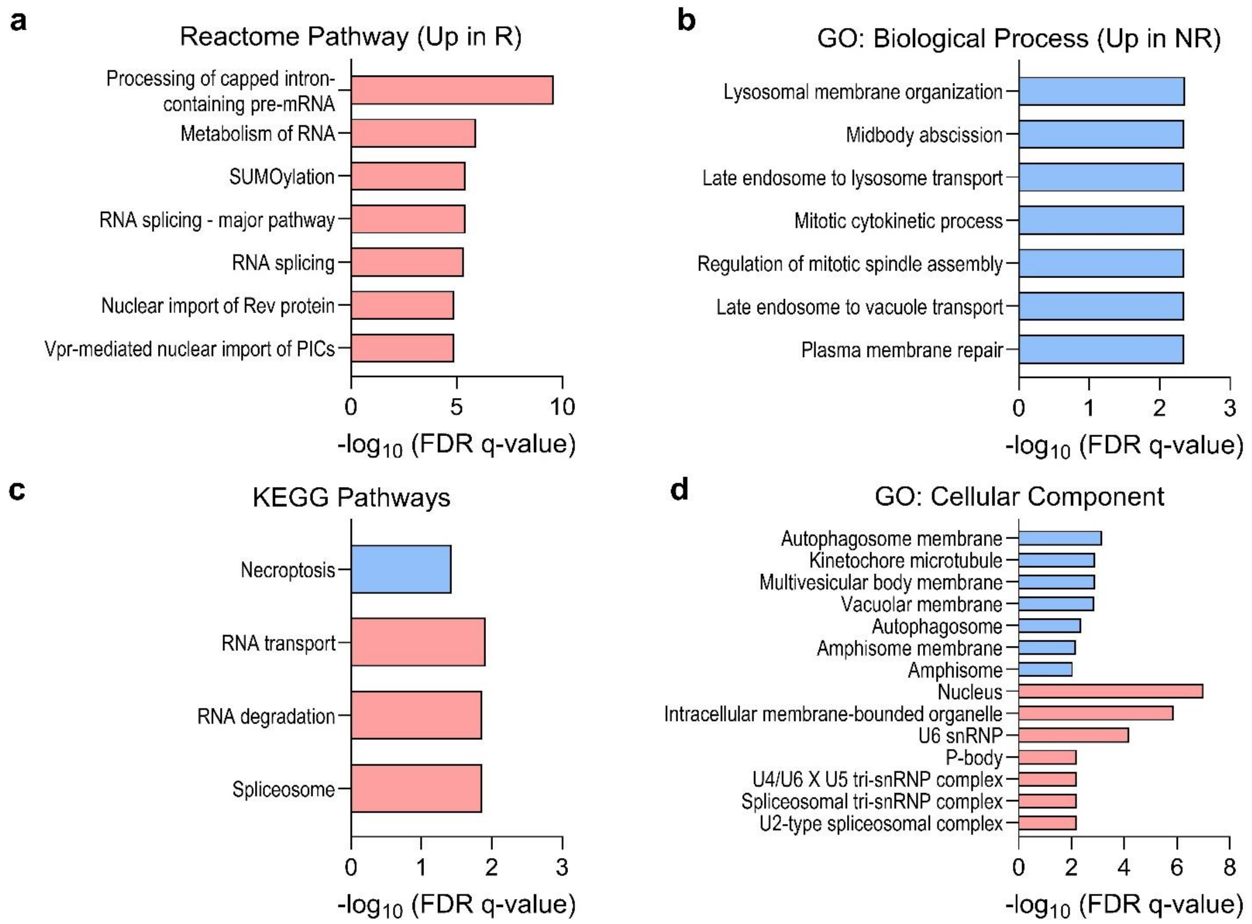


**Fig. S4. Post-LVAD responder proteomes display increased RNA processing factors and decreased autophagosome membrane proteins. a-d.** Pathway, biological processes, and cell components enriched in post-LVAD responder and non-responder proteomes; FDR-adjusted p-values were used to select pathways/processes/components that were significantly enriched. Blue color = increased in non-responders; Red color = increased in responders.

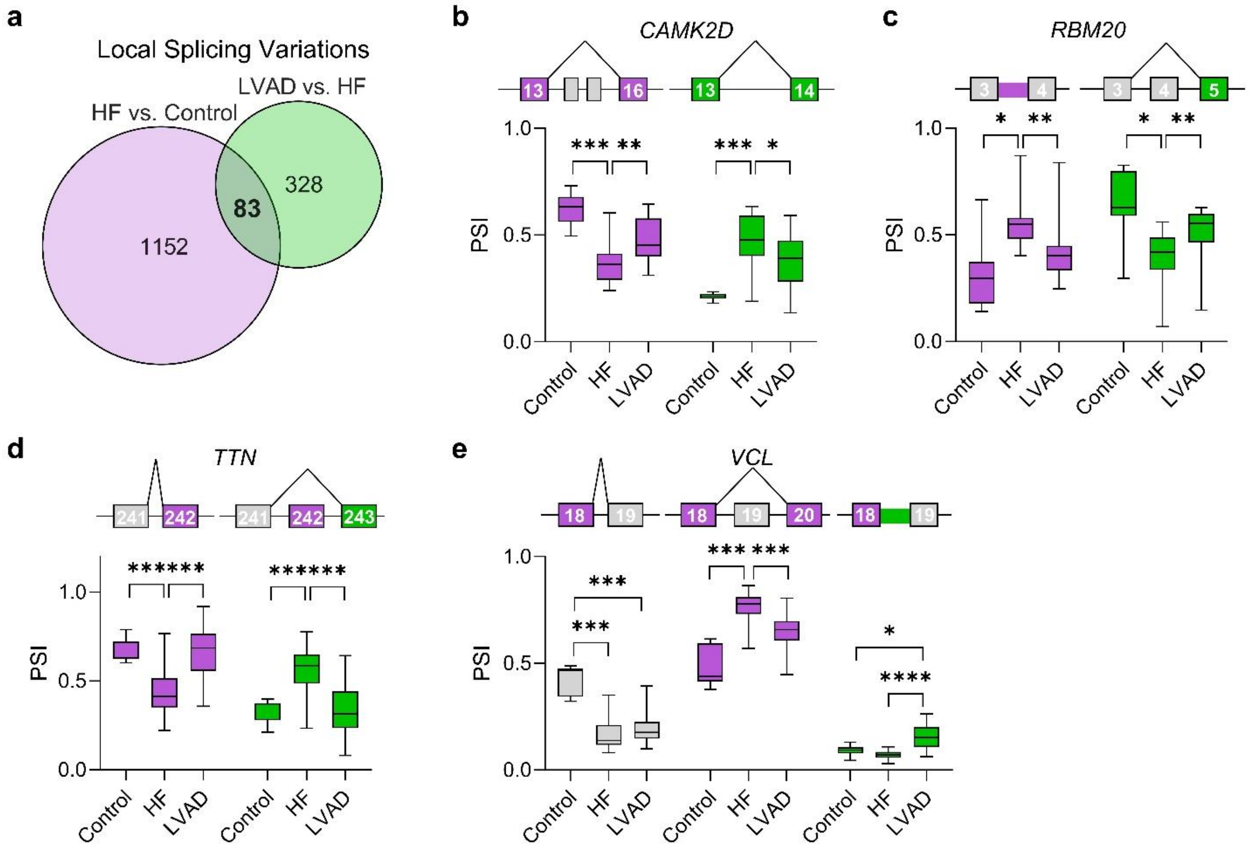


**Fig. S5. Alternative splicing changes in heart failure and with LVAD therapy. a.** Venn Diagram depicting the number of significant local splicing variations (LSVs) identified by short-read RNA-sequencing analysis between HF and non-failing controls or post-LVAD. **b-e.** Examples of differential alternative splicing between control, HF (pre-LVAD), and post-LVAD patients. *RBM20* splicing changes in HF that are reversed by LVAD include increased intron retention between exons 3 and 4 and decreased exon 4 inclusion (**b**). *TTN* exon 242 inclusion decreases in HF and is reversed by LVAD (**c**). *CAMK2D* exon 16 inclusion decreases in HF, while exon 14 inclusion increases; both events are partially reversed by LVAD therapy (**d**). *VCL* exon 19 inclusion decreases in heart failure. LVAD modestly reduces exon 19 skipping while increasing intron retention. n = 5 non-failing controls, n = 19 HF and LVAD. *p < 0.05, **p < 0.01, ***p < 0.001, ****p < 0.0001 by an independent two-sample Mann-Whitney U-test (Wilcoxon test in MAJIQ).

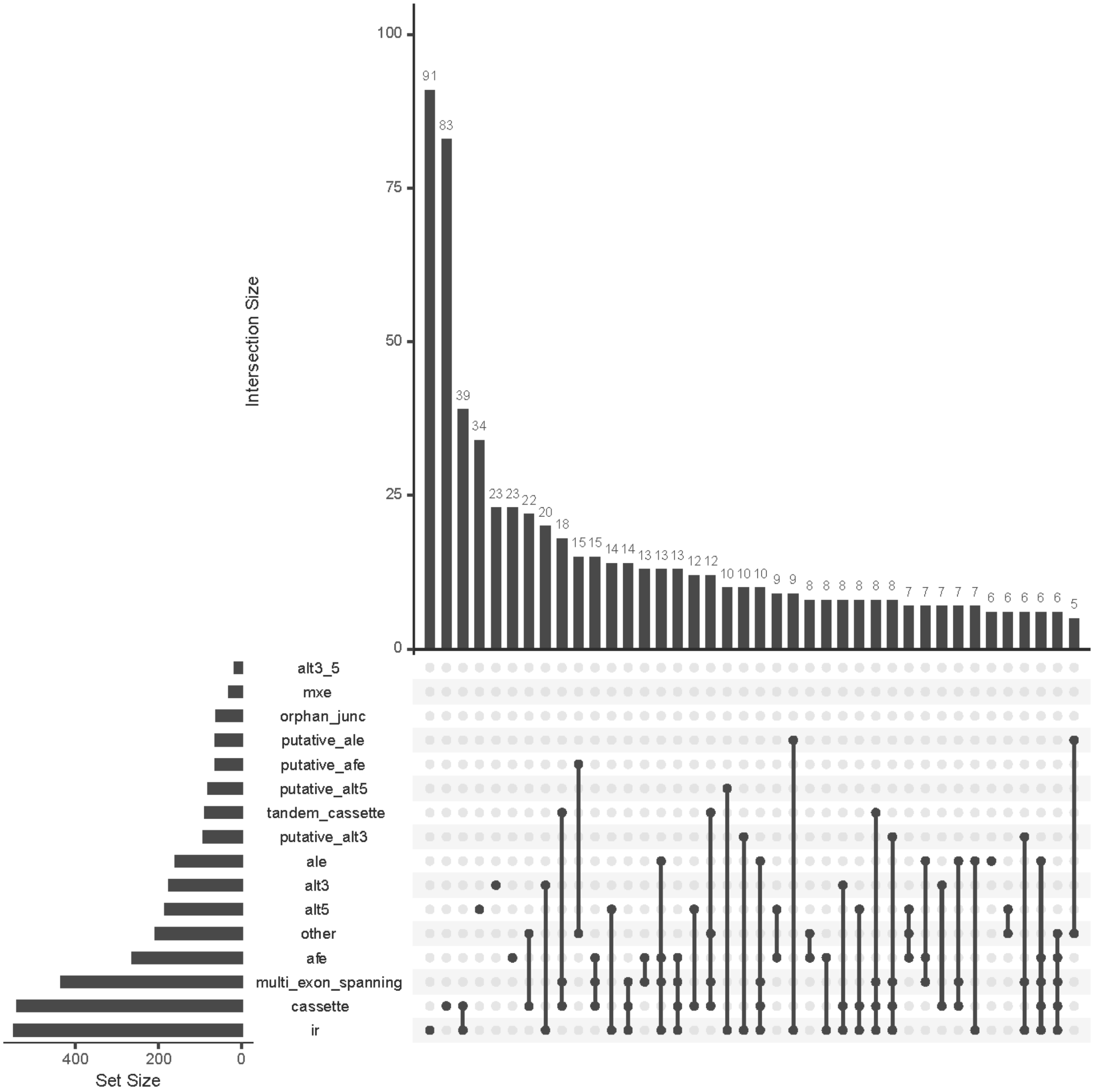


**Fig. S6. Alternative splicing changes with end-stage heart failure.** Graphical representation of the alternative splicing modules that undergo differential splicing between HF and Control. The splicing events contained within those modules are shown. Modules were defined as changing if they contained a changing junction. See methods for definitions of significant changes. alt3: alternative 3’ splice site. alt5: alternative 5’ splice site. Mxe: mutually exclusive exons. Orphan_junc: orphan junction. Ale: alternative last exon. Afe: alternative first exon. ir: intron retention.

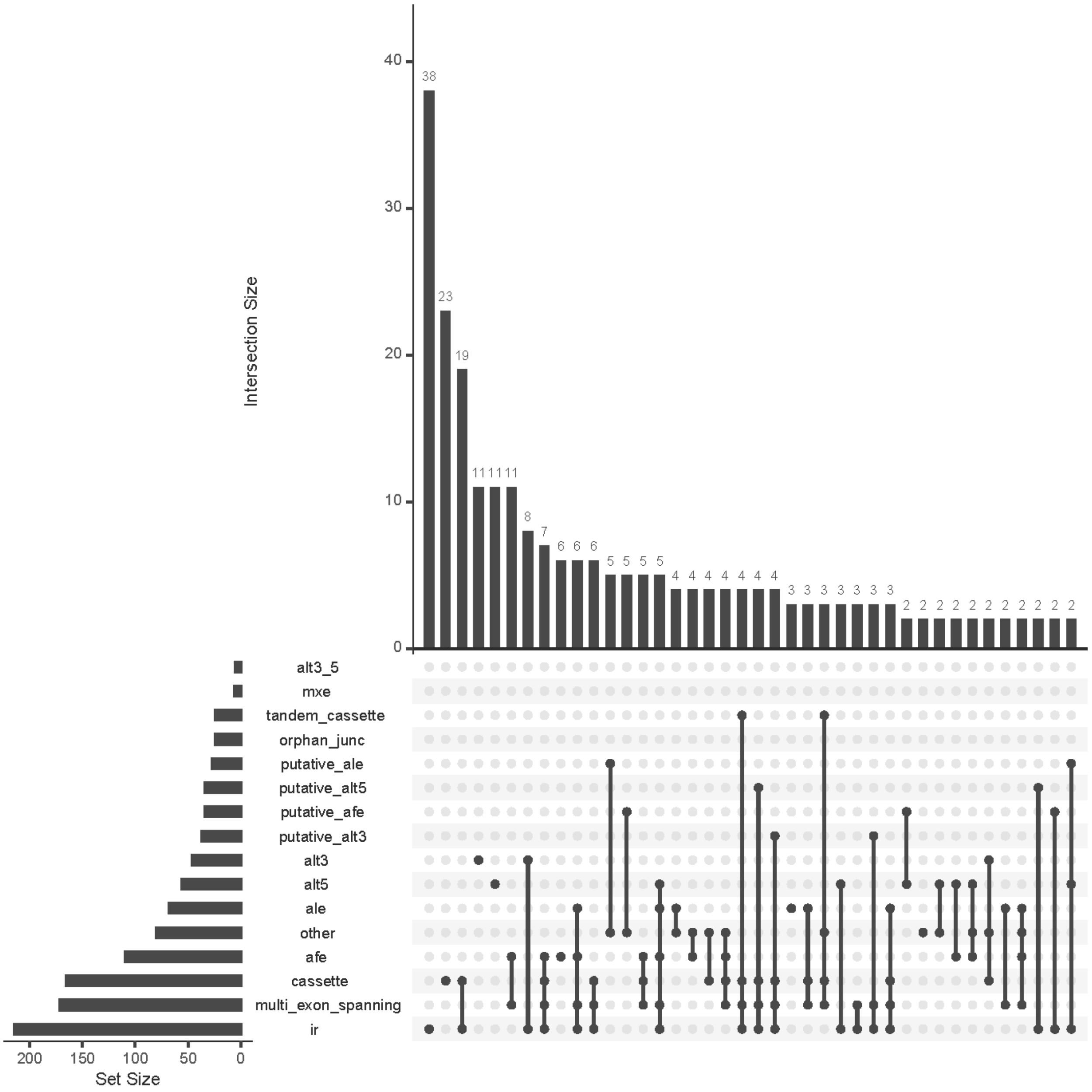


**Fig. S7. Alternative splicing changes with LVAD.** Graphical representation of the alternative splicing modules that undergo differential splicing between LVAD and HF. The splicing events contained within those modules are shown. Modules were defined as changing if they contained a changing junction. See methods for definitions of significant changes. alt3: alternative 3' splice site. alt5: alternative 5' splice site. Mxe: mutually exclusive exons. Orphan_junc: orphan junction. Ale: alternative last exon. Afe: alternative first exon. ir: intron retention.

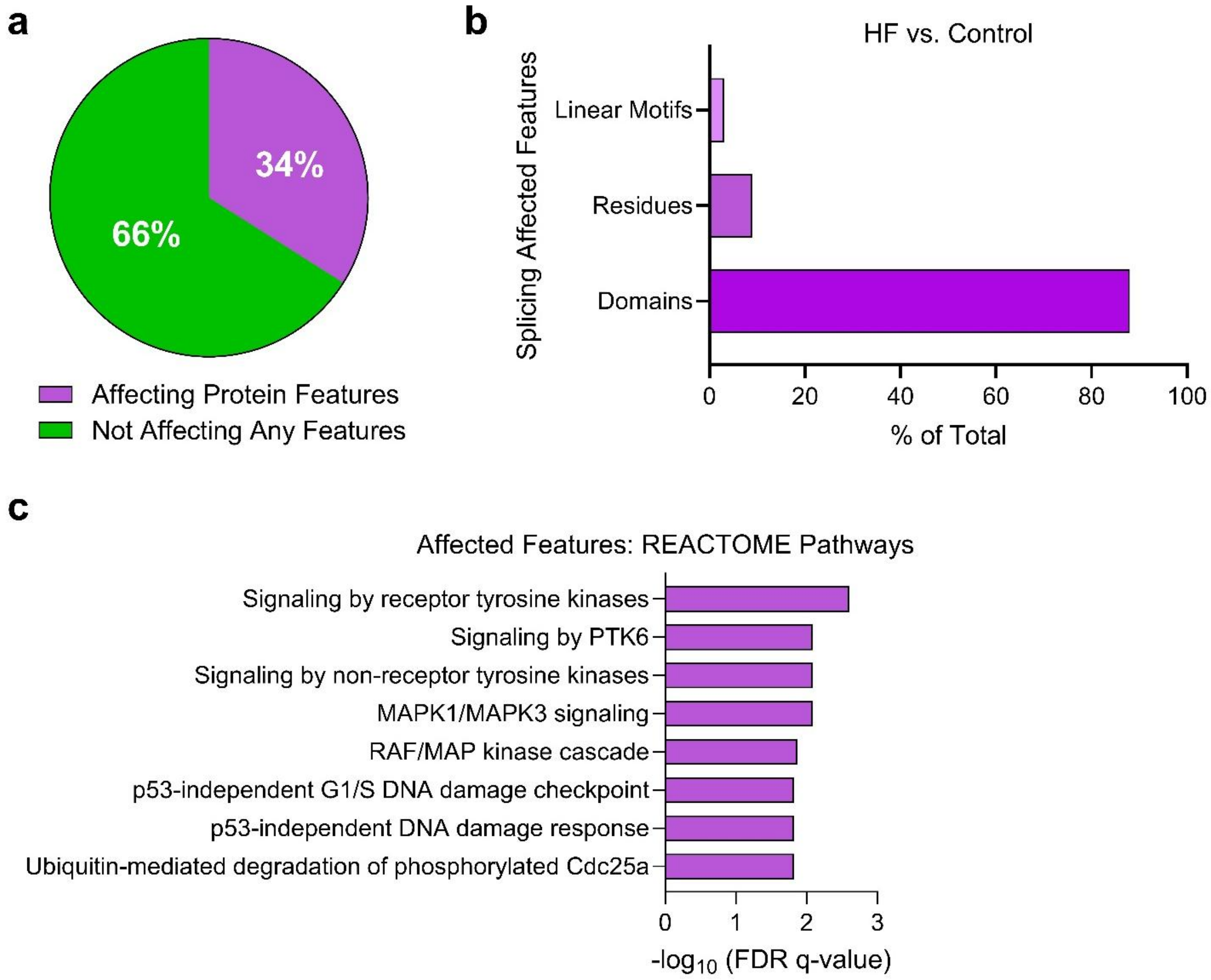


**Fig. S8. Heart failure-associated protein features affected by cassette exon alternative splicing. a.** Pie chart depicting the proportion of all HF-associated cassette exon splicing changes (vs. Control) impacting protein features. **b.** Types of protein features impacted by cassette exon alternative splicing in heart failure. **c.** REACTOME pathway over-enrichment of the pthways affected by cassette exon alternative splicing and subsequent changes in protein-protein interactions.

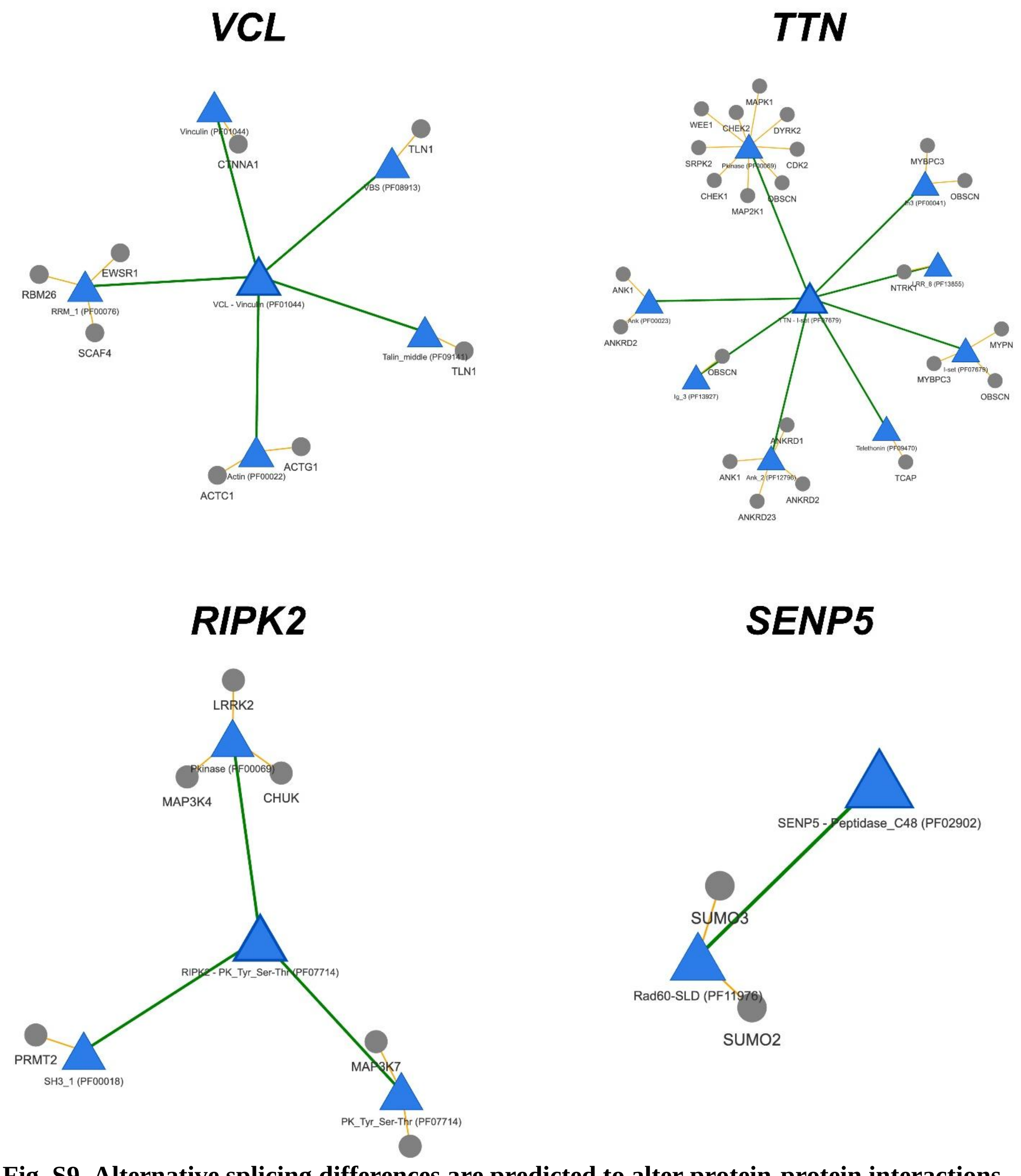


**Fig. S9. Alternative splicing differences are predicted to alter protein-protein interactions in heart failure patients.** Examples of four genes where heart-failure associated differential alternative splicing of cassette exons impacts protein domains that are known to mediate domain-domain interactions. Domains are shown as blue triangles, while proteins with predicted disrupted binding are shown as gray circles.

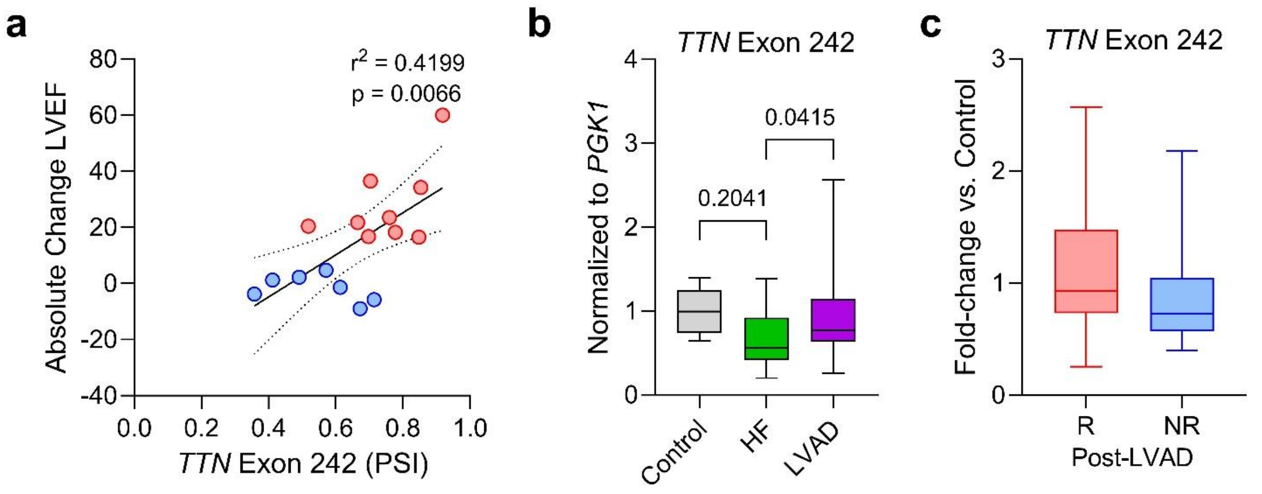


**Fig. S10. *TTN* exon 242 alternative splicing in heart failure and post-LVAD. a.** Linear regression analysis of *TTN* exon 242 proportion spliced in (PSI) post-LVAD identified by RNA-seq versus absolute change in LVEF with LVAD therapy; confidence interval = 95%. **b.** qPCR quantification of exon 242 inclusion in control, HF, and LVAD using a forward primer spanning the exon-exon junction of exons 241 and 242; one-way ANOVA with Tukey's post-hoc test for multiple independent pairwise comparisons; n = 6 control, 19 HF, 19 LVAD. **c.** Quantification of exon 242 inclusion between post-LVAD responders and non-responders from the qPCR analysis; n = 10 R, 9 NR.

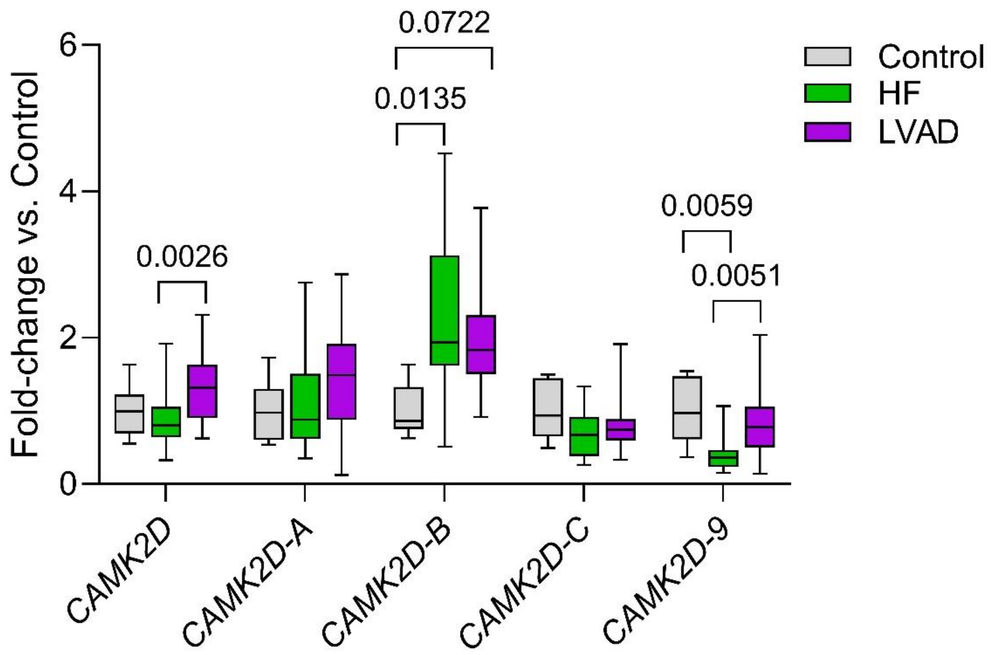


**Fig. S11. *CAMK2D* alternative isoform expression is impacted by heart failure and partially restored with mechanical circulatory support.** Expression of total *CAMK2D*, *CAMK2D-A* (Exons 13-15), *CAMK2D-B* (Exons 13-14), *CAMK2D-C* (Exons 13-17), and *CAMK2D-9* (Exons 13-16) by qPCR between non-failing controls (n = 6), heart failure/pre-LVAD (n = 19), and heart failure/post-LVAD (n = 19). For each, data were analyzed by one-way ANOVA with Tukey's post-hoc test for multiple independent pairwise comparisons.

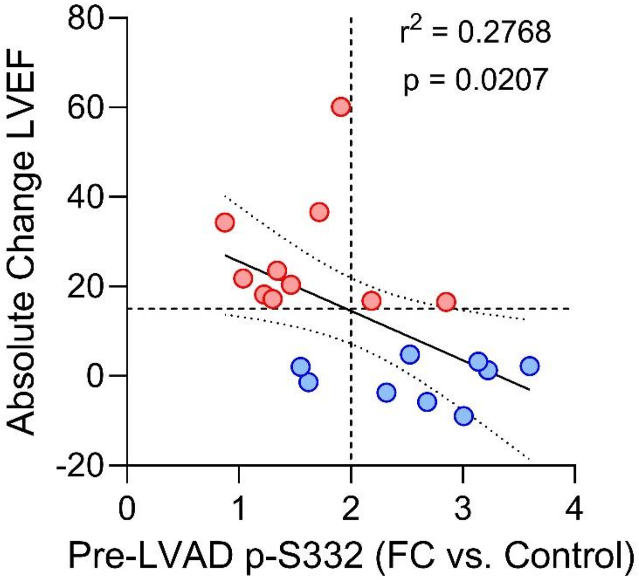


**Fig. S12. CaMKIIδ-B S332 phosphorylation in heart failure negatively correlates with functional recovery post-LVAD.** Linear regression analysis of pre-LVAD S332 phosphorylation versus absolute change in LVEF with LVAD therapy; n = 19 patients; red = R, blue = NR; y-axis line = 15, x-axis line = 2; confidence interval = 95%.

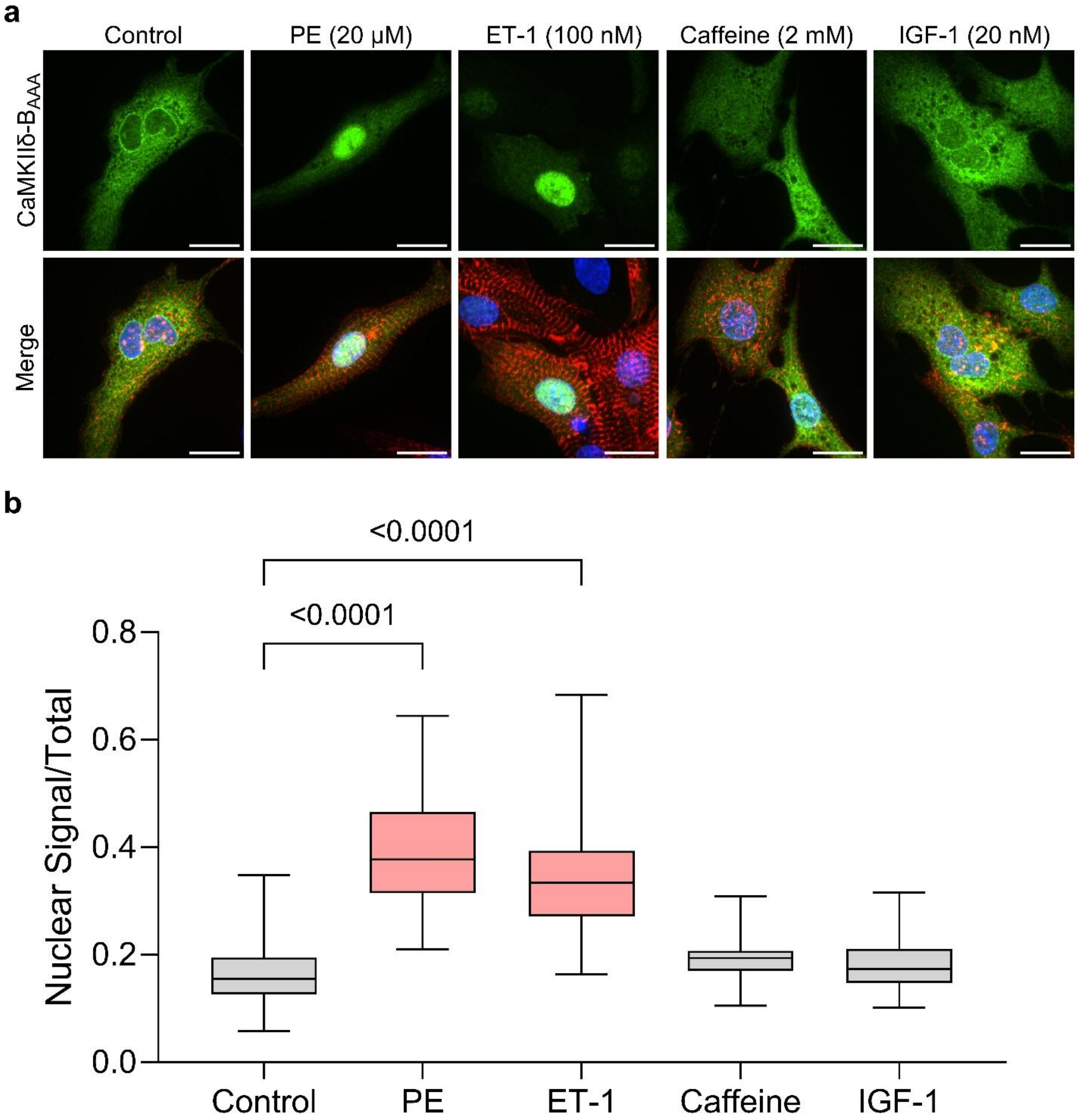


**Fig. S13. Adrenergic and endothelin receptor agonists induce CaMKIIδ-$B_{AAA}$ translocation to the nucleus. a.** Representative immunofluorescence microscopy images of NRVMs transduced with GFP-$B_{AAA}$ and treated with phenylephrine (PE), endothelin-1 (ET-1), caffeine, or insulin-like growth factor-1 (IGF-1). **b.** Quantification of nuclear GFP signal to total cellular GFP signal; one-way ANOVA with Tukey's post-hoc test for multiple independent pairwise comparisons; n = 73 Control, 85 PE, 53 ET-1, 55 Caffeine, and 45 IGF-1 from three independent biological replicates.

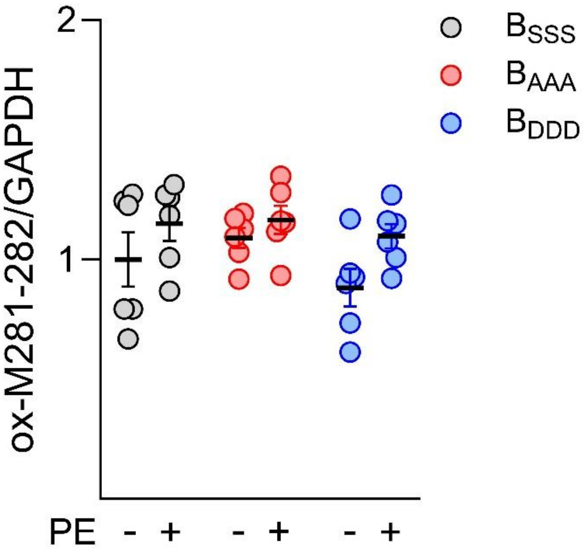


**Fig. S14. Impact of phenylephrine on CaMKIIδ-B oxidation.** Oxidation of M281/M282 GFP-CaMKIIδ-B normalized to GAPDH in NRVMs treated with PE for 24 hours. Representative western blot is depicted in Figure 4; n = 6/group.

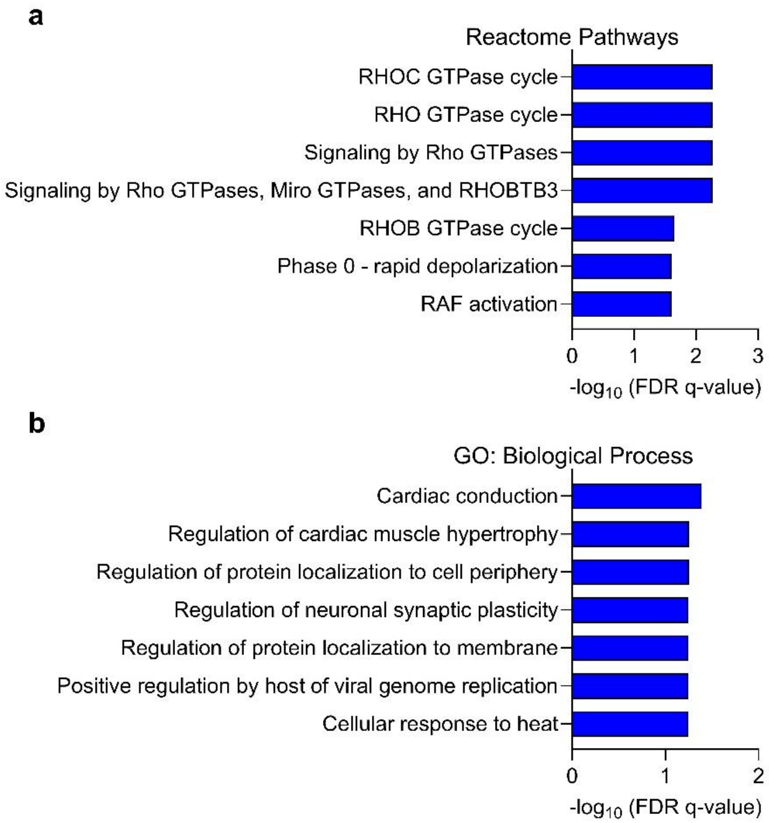


**Figure S15. Pathway enrichment for phospho-peptides correlated with p-S332-334 CaMKIIδ post-LVAD. a.** Reactome Pathway enrichment for phospho-peptides with $r^2 > 0.50$ vs. CaMKIIδ-B S332-S334 phosphorylation by linear regression. **b.** GO: Biological Process enrichment for phospho-peptides with $r^2 > 0.50$ vs. CaMKIIδ-B S332-S334 phosphorylation by linear regression.

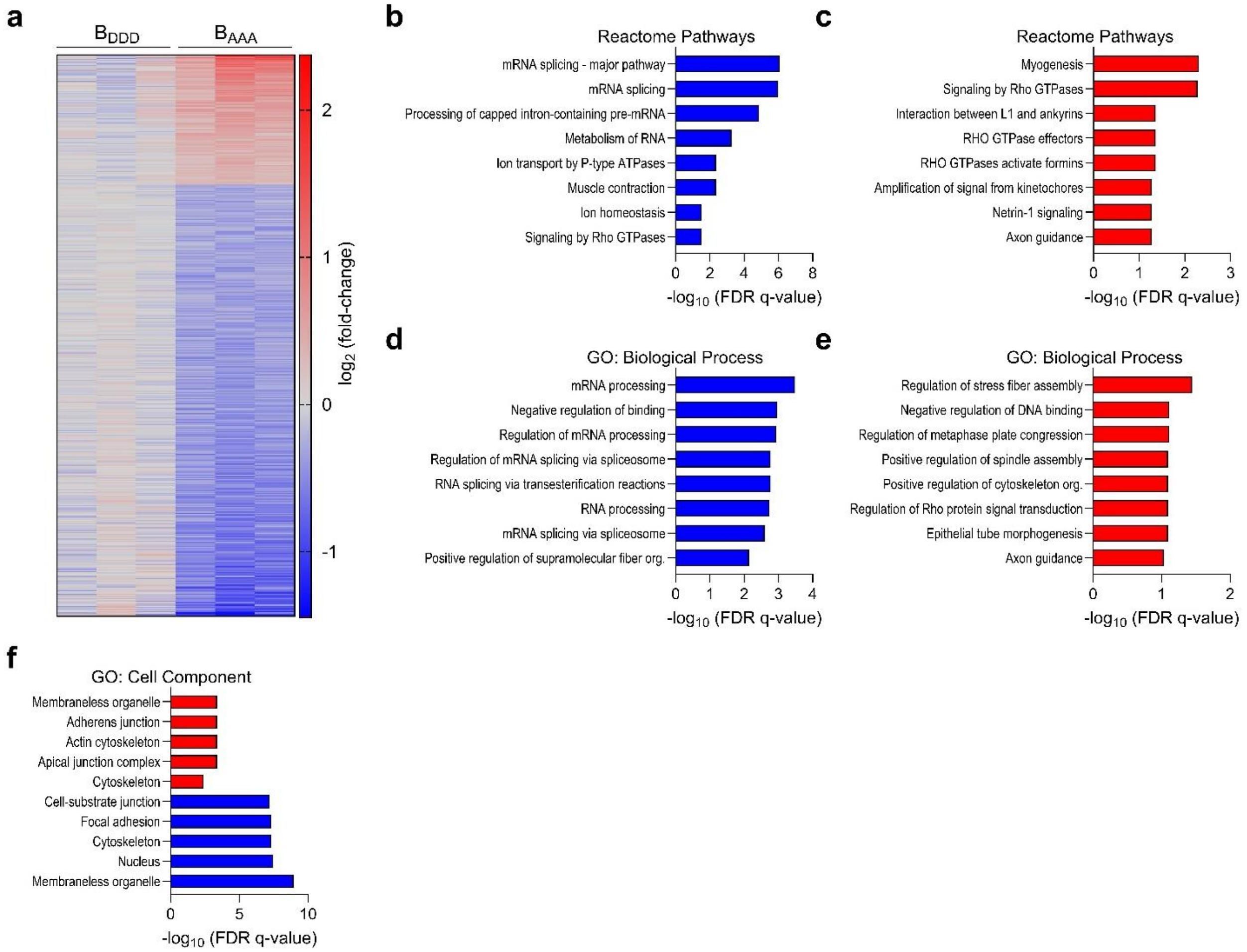


**Fig. S16. Phospho-proteome differences between B-DDD and B-AAA transduced NRVMs.** **a.** Heat map of significantly differentially phosphorylated protein residues between B-AAA and B-DDD transduced NRVMs identified by phospho-proteomics; p-value < 0.05 and $log_2$-FC < -0.3, > 0.3. **b-c.** Significantly over-enriched Reactome Pathways in the B-DDD (b) and B-AAA (c) cells. **d-e.** Significantly over-enriched GO: Biological Processes in the B-DDD (d) and B-AAA (e) cells. **f.** GO Cell Component enrichment for the differentially expressed phospho-peptides.

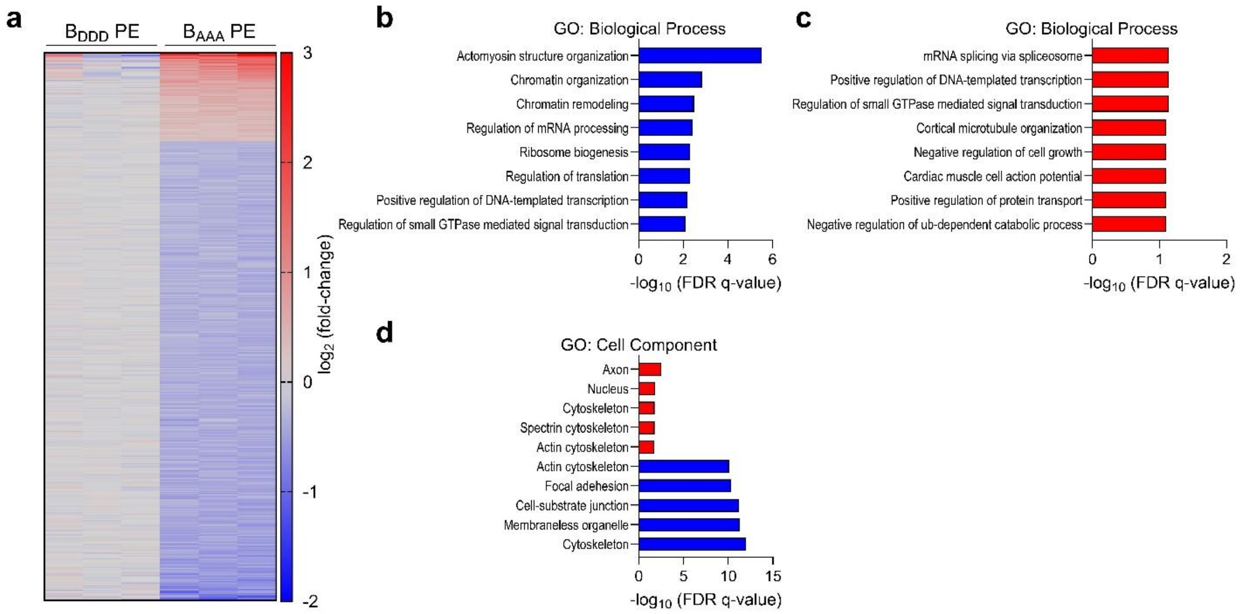


**Fig. S17. Phospho-proteome differences between B-DDD and B-AAA transduced NRVMs with adrenergic agonism. a.** Heat map of significantly differentially phosphorylated protein residues between B-AAA and B-DDD transduced NRVMs treated with PE identified by phospho-proteomics; p-value < 0.05 and $\log_2$-FC < -0.3, > 0.3. **b-c.** Significantly over-enriched GO: Biological Processes in the B-DDD/PE (b) and B-AAA/PE (c) cells. **d.** GO Cell Component enrichment for the differentially expressed phospho-peptides.

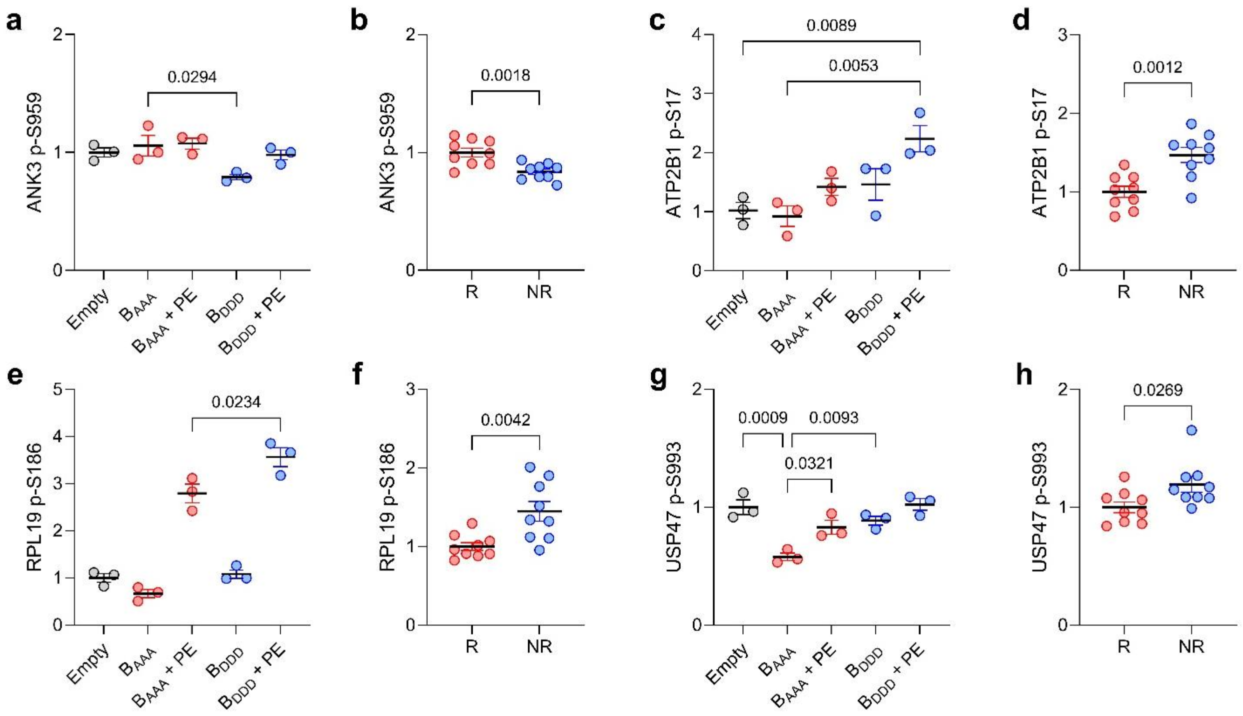


**Fig. S18. Phosphorylation differences in B$_{DDD}$-transduced NRVMs are shared in human heart failure non-responders. a-h.** Quantification of four different phospho-peptides with shared expression features in transduced NRVMs and post-LVAD R vs. NR patients; NRVMs: n = 3/group from 3 different experiments, one-way ANOVA with Tukey's post-hoc test; post-LVAD: n = 9/group, two-tailed t-test; amino acid position in humans used for rat peptides for simplicity. ANK3: ankyrin-3, ATP2B1: ATPase Plasma Membrane Ca2+ Transporting 1, RPL19: Ribosomal protein L19, USP47: Ubiquitin-specific peptidase 47.

**Table S1. Patient clinical characteristics.**

| Characteristic | Non-Failing Donors | HF/LVAD Patients | p-value |
|---|---|---|---|
| Number of patients | 6 | 19 | n.a. |
| Age (years) | 51.0 ± 16.9 | 48.1 ± 13.3 | 0.660 |
| Sex (% male) | 100 | 84.2 | 0.299 |
| Race (% white) | 83.3 | 68.4 | 0.478 |
| LVAD duration (days) | ----- | 267 ± 253 | n.a. |
| **Clinical Data** | **HF (pre-LVAD)** | **Post-LVAD** | **p-value** |
| LVEF (%) | 14.1 ± 9.8 | 27.7 ± 15.6 | 0.003 |
| LV Fractional Shortening (%) | 7.2 ± 4.2 | 13.8 ± 8.3 | 0.003 |
| LVESD (cm) | 6.4 ± 0.9 | 5.0 ± 1.6 | <0.001 |
| LVEDD (cm) | 6.9 ± 1.0 | 5.8 ± 1.5 | <0.001 |
| Serum BNP (pg/mL)* | 1993 ± 1214 | 244 ± 196 | <0.001 |
| ***Medications*** | | | |
| ACE inhibitor, n (%) | 1 (5.3) | 10 (52.6) | 0.001 |
| Aldosterone antagonist, n (%) | 15 (78.9) | 14 (73.7) | 0.703 |
| Anti-arrhythmic, n (%) | 4 (21.1) | 5 (26.3) | 0.703 |
| Beta blocker, n (%) | 3 (15.8) | 10 (52.6) | 0.017 |
| Digoxin, n (%) | 9 (47.4) | 5 (26.3) | 0.179 |
| Dobutamine, n (%) | 11 (57.9) | 1 (5.3) | 0.001 |
| Loop diuretic, n (%) | 15 (78.9) | 6 (31.6) | 0.003 |

Data are presented as the mean ± SD.
All clinical data were analyzed by two-tailed paired t-test.
Categorical data (Sex, Race, Medications) were analyzed by Chi-squared test.
*Values not available for 4 pre-LVAD and 4 post-LVAD patients

**Table S2. Clinical characteristics of responders and non-responders pre- and post-LVAD.**

| Characteristic | Responders | Non-responders | p-value |
|---|---|---|---|
| Number of patients | 10 | 9 | n.a. |
| Age (years) | 46.3 ± 15.4 | 50.0 ± 10.9 | 0.559 |
| Sex (% male) | 80.0 | 88.9 | 0.596 |
| Race (% white) | 90.0 | 44.4 | 0.033 |
| Ethnicity (% non-Hispanic) | 80.0 | 100 | 0.156 |
| LVAD duration (days) | 229 ± 268 | 308 ± 245 | 0.517 |
| | **HF (pre-LVAD)** | **HF (pre-LVAD)** | **p-value** |
| LVEF (%) | 10.4 ± 8.6 | 18.1 ± 9.9 | 0.086 |
| LVFS (%) | 6.1 ± 3.0 | 8.3 ± 5.2 | 0.272 |
| LVESD (cm) | 6.2 ± 0.8 | 6.6 ± 1.0 | 0.360 |
| LVEDD (cm) | 6.6 ± 0.8 | 7.1 ± 1.1 | 0.206 |
| LV Systolic Volume (mL) | 176 ± 53 | 227 ± 121 | 0.247 |
| LV Diastolic Volume (mL)# | 195 ± 53 | 273 ± 131 | 0.105 |
| Serum BNP (pg/mL)* | 1645 ± 1002 | 2517 ± 1404 | 0.182 |
| ***Medications*** | | | |
| ACE inhibitor, n (%) | 1 (10.0) | 0 (0.0) | 0.330 |
| Aldosterone antagonist, n (%) | 7 (70.0) | 8 (88.9) | 0.313 |
| Anti-arrhythmic, n (%) | 4 (40.0) | 0 (0.0) | 0.033 |
| Beta blocker, n (%) | 3 (30.0) | 0 (0.0) | 0.073 |
| Digoxin, n (%) | 5 (50.0) | 4 (44.4) | 0.809 |
| Dobutamine, n (%) | 5 (50.0) | 6 (66.7) | 0.463 |
| Loop diuretic, n (%) | 7 (70.0) | 8 (88.9) | 0.313 |
| | **Post-LVAD** | **Post-LVAD** | **p-value** |
| LVEF (%) | 37.0 ± 14.4 | 17.4 ± 9.4 | <0.001 |
| LVFS (%) | 17.2 ± 10.1 | 10.0 ± 3.1 | 0.055 |
| LVESD (cm) | 3.9 ± 1.0 | 6.3 ± 1.0 | <0.001 |
| LVEDD (cm) | 4.7 ± 0.8 | 7.0 ± 1.0 | <0.001 |
| LV Systolic Volume (mL)# | 50 ± 24 | 168 ± 81 | <0.001 |
| LV Diastolic Volume (mL)# | 77 ± 35 | 202 ± 93 | 0.002 |
| Serum BNP (pg/mL)* | 164 ± 82 | 335 ± 252 | 0.092 |
| ***Medications*** | | | |
| ACE inhibitor, n (%) | 7 (70.0) | 3 (33.3) | 0.110 |
| Aldosterone antagonist, n (%) | 8 (80.0) | 6 (66.7) | 0.510 |
| Anti-arrhythmic, n (%) | 2 (20.0) | 3 (33.3) | 0.510 |
| Beta blocker, n (%) | 6 (60.0) | 4 (44.4) | 0.498 |
| Digoxin, n (%) | 3 (30.0) | 2 (22.2) | 0.599 |
| Dobutamine, n (%) | 1 (10.0) | 0 (0.0) | 0.305 |
| Loop diuretic, n (%) | 2 (20.0) | 4 (44.4) | 0.252 |

Data are presented as the mean ± SD.

Categorical data were analyzed by Chi-squared test. All other data were analyzed by two-tailed unpaired t-test.

#Values not available for 1 pre-LVAD and 1 post-LVAD patient

*Values not available for 4 pre-LVAD and 4 post-LVAD patients

**Table S3. Domain interactions altered by alternative splicing in heart failure.**

| Gene Name | NCBI Gene ID | PFAM or ELM Identifier | ΔPSI (HF vs. Control) | # of Affected Interactions | Affected Binding |
|---|---|---|---|---|---|
| TTN | 7273 | PF07679 | -0.216 | 17 | ANKRD23, TCAP, CHEK1, ANKRD2, ANKRD1, MYBPC3, NTRK1, CDK2, CHEK2, OBSCN, MAPK1, WEE1, SRPK2, ANK1, MYPN, MAP2K1, DYRK2 |
| NF1 | 4763 | PF00616 | -0.181 | 15 | PHLDB2, RASAL2, KRAS, GAB2, SRGAP2, AGAP1, HRAS, RTKN, SH3RF3, RALGPS2, NRAS, AKT1, TIAM1, SH3PXD2A, PLEKHA7 |
| PTPN20 | 26095 | PF00102 | 0.183 | 10 | KIT, AATK, PTPN6, PTPN1, ERBB4, PTPN2, ERBB3, PTPN20, EGFR, ROR2 |
| VCL | 7414 | PF01044 | -0.328 | 8 | SCAF4, ACTG1, EWSR1, TLN1, ACTC1, RBM26, VCL, CTNNA1 |
| RIPK2 | 8767 | PF07714 | 0.392 | 7 | PRMT2, MAP3K4, LRRK2, CHUK, RIPK1, MAP3K7, RIPK2 |
| MLKL | 197259 | PF07714 | 0.256 | 4 | RIPK1, LRCH3, CDK1, RIPK3 |
| ZNF384 | 171017 | PF00096 | -0.152 | 4 | UBE2K, HOXC9, TRIP6, NKX2 |
| ABI2 | 10152 | PF07815 | 0.251 | 4 | NCKAP1, NCKAP1L, CYFIP2, CYFIP1 |
| NCAM1 | 4684 | PF13927 | -0.195 | 4 | NTRK1, FGFR1, OPCML, SIGLEC9 |
| ANKS3 | 124401 | PF12796 | 0.235 | 3 | ANKS6, NEKB, NEK7 |
| UIMC1 | 51720 | PF18282 | -0.159 | 3 | RAD23A, UBB, UBC |
| PLK3 | 1263 | PF00659 | -0.177 | 3 | PLK1, CHEK2, AURKA |
| SENP5 | 205564 | PF02902 | 0.283 | 2 | SUMO3, SUMO2 |
| FOXJ3 | 22887 | PF00250 | -0.174 | 2 | ZBTB2, FOXJ1 |
| ASPG | 374569 | PF12796 | 0.169 | 2 | PRKD2, SGK1 |
| FKBP7 | 51661 | PF00254 | 0.165 | 2 | SGTA, SOD1 |
| POLK | 51426 | PF11799 | 0.212 | 2 | PCNA, REV1 |
| 1601 | 1601 | LIG_AP2alpha_2 | 0.155 | 2 | AP2A1, AP2A2 |
| 65125 | 65125 | DOC_SPAK_OSR1_1 | -0.169 | 2 | WNK2, WNK1 |
| YTHDC2 | 64848 | PF01424 | 0.200 | 1 | NFX1 |
| GABBR1 | 2550 | PF00003 | -0.172 | 1 | GABBR2 |
| GALK2 | 2585 | PF00288 | 0.212 | 1 | MVD |

**Table S4. Top phospho-sites correlated with CaMKIIδ-B NLS phosphorylation in post-LVAD human hearts.** Previously identified CaMKII substrates/binding partners are denoted by ***bold italics***.

| Gene | Centralized Sequence | Position | Linear Regression ($r^2$) |
|---|---|---|---|
| CAMK2D | KSLLKKPDGVKKRKSSSSVQMMESTESSNTT | 333 | 1 |
| CAMK2D | SLLKKPDGVKKRKSSSSVQMMESTESSNTTI | 334 | 0.973692373 |
| CAMK2D | AKSLLKKPDGVKKRKSSSSVQMMESTESSNT | 332 | 0.936509652 |
| CAMK2D | SLLKKPDGVKKRKSSSSVQMMESTESSNTTI | 334 | 0.897938799 |
| DMXL1 | RASSFLDTSKDCSPSSPLKLDAREDKSSAVD | 1908 | 0.704887268 |
| RBM20 | ________MVLAAAMSQDADPSGPEQPDRVA | 8 | 0.686864212 |
| HECTD1 | TPGESSAISMGIVSVSSPDVSSVSELTNKEA | 1487 | 0.680826617 |
| SUPT7L | YWGEIPISSSQTNRSSFDLLPREFRLVEVHD | 19 | 0.663084951 |
| CAMK2A | GGKSGGNKKSDGVKESSESTNTTIEDEDTKV | 310 | 0.641926859 |
| CAMK2D | AKSLLKKPDGVKKRKSSSSVQMMESTESSNT | 332 | 0.630109621 |
| SPTBN1 | STKVSEEAESQQQWDTSKGEQVSQNGLPAEQ | 2121 | 0.624107198 |
| POLDIP3 | RQVADAREKISLKRSSPAAFINPPIGTVTPA | 127 | 0.622969865 |
| ***CACNB2*** | ***RAKQGKFYSSKSGGNSSSSLGDIVPSSRKST*** | ***204*** | ***0.619841746*** |
| MLIP | ENSHTLLSHNACNKLSHPMVAIPEHEALDSK | 976 | 0.618451902 |
| AKAP8 | EPDTKLARVDSEGDFSENDDAAGDFRSGDEE | 315 | 0.617454573 |
| ***BAZ1A*** | ***SLQESESKRRCRKRQSPEPSPVTLGRRSSGR*** | ***1413*** | ***0.610216459*** |
| TPR | KRKGAILSEEELAAMSPTAAAVAKIVKPGMK | 379 | 0.607747849 |
| ATXN3 | QLSMQGSSRNISQDMTQTSGTNLTSEELRKR | 269 | 0.607562714 |
| DDX55 | EKKKKMNEKRKREEGSDIEDEDMEELLNDTR | 513 | 0.607445905 |
| ***CRKL*** | ***YLDTTTLIEPAPRYPSPPMGSVSAPNLPTAE*** | ***107*** | ***0.604695177*** |
| ***SETD1B*** | ***KRLRPSTSVDEEDEESERERDRDMADTPCEL*** | ***994*** | ***0.602277635*** |
| ***LDB3*** | ***TTSISKQTLPRGGPAYTPAGPQVPPLARGTV*** | ***458*** | ***0.60180993*** |
| CAMK2A | NKKSDGVKESSESTNTTIEDEDTKVRKQEII | 316 | 0.599574257 |
| FAM160B2 | SGFQTPAKPRLAPATSYDGKTAVTEIVNSFL | 526 | 0.599342867 |
| ***RPL19*** | ***EERLQAKKEEIIKTLSKEEETKK________*** | ***186*** | ***0.595835997*** |
| RALY | ARTRDDGDEEGLLTHSEEELEHSQDTDADDG | 272 | 0.589595084 |
| R3HDM2 | STDSELKSLEPRPWSSTDSDGSVRSMRPPVT | 344 | 0.589494994 |
| HELZ | KQVDLESNPQNRSPESRPSVVYPSTKFPRKD | 1321 | 0.586938791 |
| THRAP3 | NQGDEAKEQTFSGGTSQDTKASESSKPWPDA | 211 | 0.584987684 |
| THRAP3 | DNQGDEAKEQTFSGGTSQDTKASESSKPWPD | 210 | 0.584987684 |
| MSL3 | FLHLEKKTPVHSRSSSPIPLTPSKEGSAVFA | 367 | 0.581408118 |
| MFAP1 | RYVSGKRPDYAPMESSDEEDEEFQFIKKAKE | 53 | 0.578349581 |
| MFAP1 | KRYVSGKRPDYAPMESSDEEDEEFQFIKKAK | 52 | 0.578349581 |
| MAP1B | PTPVVKQTKLKQRADSRESLKPAAKPLPSKS | 541 | 0.577619534 |
| ***GPRIN3*** | ***TATSLSLPSDPMGDSSPGSGKKTPSRSVKAS*** | ***616*** | ***0.577131044*** |
| SRCIN1 | VVTSKKDSAFIKKAESEELEVQKPQVKLRRA | 1077 | 0.574232871 |

| | | | |
|---|---|---|---|
| CTF1 | _________MSRREGSLEDPQTDSSVSLLPH | 7 | 0.572842571 |
| MTM1 | ___MASASTSKYNSHSLENESIKRTSRDGVN | 13 | 0.572148288 |
| SON | VEYQMKSVLKSVESTSPEPSKIMLVEPPVAK | 283 | 0.568448975 |
| THRAP3 | SFSITREAQVNVRMDSFDEDLARPSGLLAQE | 575 | 0.568203487 |
| AAK1 | SDVTHSAVFGVPASKSTQLLQAAAAEASLNK | 652 | 0.567544905 |
| PEX1 | DQLRADISIIKGRYRSQSGEDESMNQPGPIK | 1152 | 0.567040217 |
| TMEM115 | LNERLKRVEDQSIWPSMDDDEEESGAKVDSP | 306 | 0.565592629 |
| TRAPPC10 | GIICRNVHMLLRRQESSSSLEMPSGVALEEG | 708 | 0.563717452 |
| SVIL | EYGETESKRALTGRDSGMEKYGSFEEAEASY | 961 | 0.563459128 |
| MCAM | VKSDKLPEEMGLLQGSSGDKRAPGDQGEKYI | 627 | 0.561923046 |
| ZNF830 | LTIKELQKKEEENADSDDEGELQDLLSQDWR | 351 | 0.561200025 |
| LMO7 | RETSVRIYQYRRPVDSYDIPKTEEASSGFLP | 1348 | 0.560801751 |
| MYLK3 | LLQKYIAQRKWKKHFYVVTAANRLRKFPTSP | 804 | 0.559909349 |
| PLEKHA5 | KGSHFPVGVVPPRAKSPTPESSTIASYVTLR | 913 | 0.558966114 |
| SMCR8 | IEVLGTEKSTSVLSKSDSQASLTVPLSPQVV | 487 | 0.558797172 |
| ABLIM2 | SSESIISVPASSTSGSPSRVIYAKLGGEILD | 299 | 0.556446182 |
| ***MEF2A*** | ***AGSSPVGNGFVNSRASPNLIGATGANSLGKV*** | ***239*** | ***0.554570413*** |
| FAM169A | SLTASINKLESTARPSESSEEFLEEEPEQRG | 316 | 0.553822683 |
| PKN2 | MRRLLRRNPERRLGASEKDAEDVKKHPFFRL | 855 | 0.551277215 |
| GATAD2B | VDMSARRSEPERGRLTPSPDIIVLSDNEASS | 120 | 0.550867142 |
| NUCKS1 | NKRRSGKNSQEDSEDSEDKDVKTKKDDSHSA | 61 | 0.550395642 |
| NUCKS1 | EAKNKRRSGKNSQEDSEDSEDKDVKTKKDDS | 58 | 0.550395642 |
| NUCKS1 | SSPREAKNKRRSGKNSQEDSEDSEDKDVKTK | 54 | 0.550395642 |
| MYCBP2 | ILPARVKAIPRRRVNSGDTEVGSSLLRHPSP | 3505 | 0.549486856 |
| DSP | IVDPVSNLRLPVEEAYKRGLVGIEFKEKLLS | 2316 | 0.547441819 |
| ***YWHAE*** | ***KNVIGARRASWRIISSIEQKEENKGGEDKLK*** | ***65*** | ***0.545649413*** |
| ***CACNB2*** | ***RAKQGKFYSSKSGGNSSSSLGDIVPSSRKST*** | ***204*** | ***0.540157067*** |
| CD2BP2 | HSLDSDEEEDDDDGGSSKYDILASEDVEGQE | 60 | 0.538827142 |
| ZC3H13 | NSKTNQSKKKGPRTPSPPPPIPEDIALGKKY | 265 | 0.535398754 |
| ZC3H13 | QRNSKTNQSKKKGPRTPSPPPPIPEDIALGK | 263 | 0.535398754 |
| ***CLASP2*** | ***STVSSGVQRVLVNSASAQKRSKIPRSQGCSR*** | ***722*** | ***0.535122544*** |
| ASPSCR1 | AADVLVARYMSRAAGSPSPLPAPDPAPKSEP | 500 | 0.534568173 |
| GATAD2B | MSARRSEPERGRLTPSPDIIVLSDNEASSPR | 122 | 0.533956437 |
| VAPB | ASKTETPIVSKSLSSSLDDTEVKKVMEECKR | 160 | 0.533269237 |
| RABGAP1L | DNSTKHEEKPQLKIVSNGDEQLEKAMEEILR | 49 | 0.532941276 |
| TNKS1BP1 | QDRVVGKPAQLGTQRSQEADVQDWEFRKRDS | 836 | 0.531958044 |
| ARHGEF11 | LHQSASSSTSSLSTRSLENPTPPFTPKMGRR | 703 | 0.529432663 |
| ADPRHL1 | KLRASVPEPRTQAGESQERPLTQADLGRQQS | 1367 | 0.526343373 |
| CMYA5 | PEDVLSQGKESFEHISENEFASEAEQSTPAE | 3462 | 0.525014008 |
| FTH1 | NAMECALHLEKNVNQSLLELHKLATDKNDPH | 114 | 0.524250975 |

| | | | |
|---|---|---|---|
| ABL2 | ENKENIEGAQDATENSASSLAPGFIRGAQAS | 566 | 0.522716329 |
| DIDO1 | PRQEAIPDLEDSPPVSDSEEQQESARAVPEK | 809 | 0.522348789 |
| DIDO1 | KKTAPRQEAIPDLEDSPPVSDSEEQQESARA | 805 | 0.522348789 |
| MACF1 | RYLEFSDRKDLHHQGSKSDDKLCGTLKSEIA | 3252 | 0.522139257 |
| ***MARK3*** | ***EYERNGRYEGSSRNVSAEQKDENKEAKPRSL*** | ***666*** | ***0.521320513*** |
| THRAP3 | PALKSPLQSVVVRRRSPRPSPVPKPSPPLSS | 264 | 0.520861973 |
| PBDC1 | GEEENTKNGGEKGADSGEEKEEGINREDKTD | 197 | 0.520200859 |
| CLCN7 | ______________MSSVELDDELLDPDMDP | 2 | 0.51608108 |
| HTATSF1 | DDDSNEKLFDEEEDSSEKLFDDSDERGTLGG | 714 | 0.515209212 |
| ***TTN*** | ***PEVPPPWKQEGYVASSSEAEMRETTLTTSTQ*** | ***354*** | ***0.515019811*** |
| TMEM230 | ATGIPSSKVKYSRLSSTDDGYIDLQFKKTPP | 24 | 0.514852534 |
| R3HDM1 | DNQIRVPLQDGRRSKSIEEREEEYQRVRERI | 283 | 0.513969846 |
| LIPE | DSEELSSLIKSNGQRSLELWPRPQQAPRSRS | 329 | 0.513434875 |
| TRAFD1 | SPELPRRRVRHQGDLSSGYLDDTKQETANGP | 430 | 0.510993072 |
| TNS1 | PLRRRAASDGQYENQSPEATSPRSPGVRSPV | 907 | 0.510728955 |
| CD2BP2 | SGGPGSRFKGKHSLDSDEEEDDDDGGSSKYD | 49 | 0.510446006 |
| NUDCD3 | EKVQPPGPVKEMAHGSQEAEAPGAVAGAAEV | 146 | 0.510437298 |
| ZSCAN31 | KEMEHLGDSKLQRDVSLDSKYRETCKRDSKA | 52 | 0.510187515 |
| PLEKHA6 | SGSMREKRRSLQLPASPAPDPSPRPAYKVVR | 972 | 0.508540804 |
| RALY | GLLTHSEEELEHSQDTDADDGALQ_______ | 282 | 0.507380753 |
| ***SYNPO*** | ***LIDKVSTPATTTSTFSREATLIPSSRPPASD*** | ***171*** | ***0.506779742*** |
| PQLC1 | ANELNARRRSFTAADSKDEEVKVAPRRSFLV | 116 | 0.506131344 |
| TACC2 | KAQEGESTLEIRKMGSCDGEGLLTSPDQPRG | 749 | 0.505483992 |
| SMCR8 | KSTSVLSKSDSQASLTVPLSPQVVRSKAVSH | 494 | 0.505394614 |
| SLC16A1 | EQKANEQKKESKEEETSIDVAGKPNEVTKAA | 466 | 0.505201381 |
| FAM192A | KELKEYRNNLKKVGISQENKKEVEKKLTVKP | 125 | 0.504471143 |
| TCEB3 | PSCTSPHQMYVDHYRSLEEDQEPIVSHQKPG | 222 | 0.502869281 |
| ARHGEF17 | SGTLQSQASRSTISSSFGNEETPSSKEATAE | 1663 | 0.502803558 |
| LMOD2 | KKVQTVRSRPLSPVATPPPPPPPPPPPPPSS | 420 | 0.502556408 |
| LMOD2 | PKLPKKVQTVRSRPLSPVATPPPPPPPPPPP | 416 | 0.502556408 |
| CAMK2A | GKSGGNKKSDGVKESSESTNTTIEDEDTKVR | 311 | 0.502371247 |
| DBN1 | SHRRMAPTPIPTRSPSDSSTASTPVAEQIER | 339 | 0.502136312 |
| HMBS | IAMSTTGDKILDTALSKIGEKSLFTKELEHA | 52 | 0.501731279 |
| VAPB | TASKTETPIVSKSLSSSLDDTEVKKVMEECK | 159 | 0.501356068 |
| CAMK2A | KKSDGVKESSESTNTTIEDEDTKVRKQEIIK | 317 | 0.500802398 |
| MYO18B | YCHFGDGEVLAVQRKSTERLEPASSPLASRS | 2217 | 0.500577779 |

**Table S5. Identification of likely pathogenic variants in human heart failure samples.**

| Group | Sample ID | DCM Mutation | Gene | Chromosome | Position | Protein Change |
|---|---|---|---|---|---|---|
| Responder | C0048 | Yes | *TTN* | 2 | 178584905 | p.Leu21579ValfsTer12 |
| Responder | C0052 | Yes | *BAG3* | 10 | 119676820 | p.Leu423LysfsTer14 |
| Responder | C0045 | | | | | |
| Responder | C0068 | | | | | |
| Responder | C0055 | Yes | *LMNA* | 1 | 156135967 | p.Arg335Trp |
| Responder | C0095 | | | | | |
| Responder | C0114 | | | | | |
| Responder | C0156 | | | | | |
| Responder | C0129 | | | | | |
| Responder | C0186 | | | | | |
| Non-responder | C0044 | Yes | *TTN* | 2 | 178561235 | p.Trp28299* |
| Non-responder | C0074 | | | | | |
| Non-responder | C0082 | | | | | |
| Non-responder | C0134 | | | | | |
| Non-responder | C0155 | | | | | |
| Non-responder | C0164 | | | | | |
| Non-responder | C0154 | | | | | |
| Non-responder | C0152 | | | | | |
| Non-responder | C0219 | | | | | |

**Table S6. Primers for quantitative real-time PCR.**

| Target Gene | Primer Sequences (5'-3') |
|---|---|
| *PGK1* (Housekeeping) | Forward: GCCAAGTCGGTAGTCCTTATG<br>Reverse: CCCAGCAGAGATTTGAGTTCTA |
| *CAMK2D* (Total) | Forward: AGCACCCTAATATTGTGCGAC<br>Reverse: CACTGTAGTATTCTCTTGCCA |
| $CAMK2D_A$ | Forward: CAACAAAGCCAACGTGGTAAC<br>Reverse: TTCCATCAGGGTTGTGGATTAC |
| $CAMK2D_C$ | Forward: ACCAGATGGAGTAAAGGAGTCA<br>Reverse: AGCTTCGATCAGTTGTTCAGTG |
| $CAMK2D_B$ | Forward: CGAGTGTTCAGATGATGGAGTCA<br>Reverse: AGCTTCGATCAGTTGTTCAGTG |
| $CAMK2D_9$ | Forward: ATGGAGTAAAGGAGCCCCAA<br>Reverse: GATAATCTCTTGCTTTCGTGC |
| *TTN-exon242-243* | Forward: GGGAAGTCCAACTAACAGCAA<br>Reverse: TGGTCCTCAAGAGGCTTAGT |

**Table S7. Primers for RT-PCR and acrylamide gel visualization.**

| Primer | Sequence (5'-3') |
|---|---|
| Human-CAMK2D-Exon13-F | GCAGCCAAGAGTTTGTTGAA |
| Human-CAMK2D-Exon17-R | CTACATTGACATCTTCATCCTC |
| Human-PGK1-F | GCCAAGTCGGTAGTCCTTATG |
| Human-PGK1-R | CCCAGCAGAGATTTGAGTTCTA |

**Data S1. (separate file)**
Clinical information for all paired pre- and post-LVAD samples in biobank

**Data S2. (separate file)**
Clinical information for non-failing controls and LVAD responders and non-responders

**Data S3. (separate file)**
Differential gene expression analysis (RNA-seq) for Control, Heart Failure, LVAD

**Data S4. (separate file)**
Differential gene expression analysis (RNA-seq) for Non-responders vs. Responders

**Data S5. (separate file)**
Splicing affected protein domains in heart failure

**Data S6. (separate file)**
Phospho-proteomics data from post-LVAD Non-responders vs. Responders

**Data S7. (separate file)**
Quantitative proteomics data from post-LVAD Non-responders vs. Responders

**Data S8. (separate file)**
Phospho-proteomics data from NRVMs with Ad-CaMKIIδ-B (AAA vs. DDD) +/- PE
